# From forest decline to salvage logging: cascading impacts on saproxylic beetle diversity

**DOI:** 10.1101/2025.11.28.690817

**Authors:** Jérémy Cours, Aurélien Sallé, Laurent Larrieu, Carlos Lopez-Vaamonde, Jörg Müller, Guilhem Parmain, Simon Thorn, Alain Roques, Christophe Bouget

## Abstract

Understanding the cascading effects of forest decline on saproxylic communities is fundamental for optimizing the management of disturbed forests toward biodiversity conservation objectives. We postulated that the nature and intensity of cascading pathways would vary along decline gradients, encompassing stages from stand decline to mortality and subsequent salvage logging, as mediated by shifts in habitat conditions and resource availability.

This study was conducted across three representative European forest contexts: fir forests in the French Pyrenees, spruce forests in the Bavarian Alps, and oak forests in the lowlands of the Loire Valley. Within these systems, we assessed how the taxonomic and functional α-diversity of saproxylic beetles responded to variations in both the diversity and density of deadwood and tree-related microhabitats (TreMs).

Our analyses identified key cascading effects of stand decline and mortality that were shaped by the considered beetle guild and by the forest type, reflecting tree species–specific disturbance legacies. Stand decline and mortality produced distinct responses within saproxylic beetle assemblages, as different successional guilds preferentially utilized either dying or dead trees. The overall influence of decline processes was positive in conifer-dominated forests.

TreMs played a central role in mediating cascading processes structuring saproxylic beetle communities throughout the forest decline continuum. The increase in TreM heterogeneity associated with stand decline or mortality enhanced saproxylic diversity, with exposed wood and trunk injuries identified as particularly influential microhabitats. Snags and large deadwood elements, especially in spruce forests, and deadwood diversity further contributed to sustaining high levels of beetle diversity. Conversely, salvage logging exerted detrimental effects on numerous guilds, primarily through reductions in TreM diversity, decreased TreM trait dispersion, and the depletion of saproxylic TreMs.

Given the pronounced context dependency of the processes driving these cascading community dynamics, and considering the increasing frequency, severity, and spatial extent of forest disturbances and global forest decline, it is imperative to integrate this complexity into management and conservation frameworks. Addressing these mechanisms with greater precision will be critical for maintaining functional biodiversity within rapidly changing forest ecosystems.

## 1. Introduction

Natural disturbances are major drivers of forest ecosystem composition, dynamics, and processes. They generate landscape heterogeneity, shape microenvironmental conditions, and influence forest communities (Pickett & White 1985; Viljur et al. 2022; Cours et al. 2022, 2023). Climate change is altering these disturbance regimes – often intensifying, prolonging, or expanding them beyond historical ranges (Turner 2010; Seidl et al. 2017). In Europe, for example, droughts are becoming more severe, frequent, and widespread (Samaniego et al. 2018), increasingly exceeding the physiological tolerance of many tree species and resulting in widespread mortality. Such episodes can shifts forest species composition, transform habitat structure, and disrupt ecosystem functions, potentially driving forests toward novel states (Johnstone et al. 2016; Batllori et al. 2020; Senf & Seidl 2021).

Severe disturbances affect tree condition, at multiple spatial scales (Allen et al. 2015; Senf et al. 2018, 2020; McDowell et al. 2020), initiating progressive loss of vigor and eventually mortality – collectively described as stand decline or dieback. These processes abruptly or gradually increase deadwood quantities and alter tree-related microhabitats (TreMs) profiles (Zemlerová et al. 2023; Bouget et al. 2024; Chowdhury et al. 2024; Bouget & Cours 2025). Disturbances legacies such as trunk cavities or perched crown deadwood modify microclimates and create distinct environmental conditions (Franklin et al. 2000; Swanson et al. 2011; Johnstone et al. 2016; Cours et al. 2023). For instance, increasing crown dieback during decline affects forest microclimates (Swanson et al. 2011; Sallé et al. 2021; Cours et al. 2023; Le Souchu et al. 2024a).

These changes can profoundly restructure forest communities, negatively affecting some trophic or habitat guilds while creating opportunities for others (Thorn et al. 2017; Kozák et al. 2021; Cours et al. 2023). Numerous studies have shown that saproxylic beetle taxonomic and functional α-diversity increases with forest stand decline (Wermelinger et al. 2002; Beudert et al. 2015; Cours et al. 2021; Viljur et al. 2022), though these responses can be spatial scale-dependent (Cours et al. 2022), non-linear (Viljur et al. 2022), and transient (Wermelinger et al. 2025). Moreover, declining and dead trees can support distinct beetle guilds along the decline-mortality succession (Le Souchu et al. 2024a).

Forest responses to disturbance also vary with local context. Different natural disturbance types, forest structures, and tree species yield contrasting disturbance legacies (Bouget et al. 2024; Bouget & Cours 2025), leading to diverse biodiversity outcomes (Turner & Seidl 2023; Cours et al. 2023). Drought, for example, generate distinct disturbance legacies compared to insect outbreaks or windstorms. Predicting biodiversity responses to future disturbance scenarios therefore requires better understanding of these interactive effects (Catford et al. 2022).

Salvage and sanitary logging are frequently implemented during stand decline to recover economic value and reduce pest risk (Lindenmayer et al. 2008). However, these interventions remove natural legacies shortly after their formation, functioning as compounded disturbance (Cours et al. 2023; Bouget et al. 2024) and often depleting deadwood-dependent biota (Thorn et al. 2018; Georgiev et al. 2022). At the same time, salvage logging can promote guilds associated with open habitats, such as pollinators (Thorn et al. 2018; Cours et al. 2021; Wermelinger et al. 2025).

Arthropods represent a major components of forest biodiversity and contribute to key ecosystem functions including nutrient cycling, trophic webs, and pollination (Wermelinger 2021). Ongoing global declines in arthropod populations (Seibold et al. 2019; Harvey et al. 2023) underscore the need to understand their sensitivity to environmental changes. Among forest arthropods, saproxylic beetles – species dependant on dead or decaying wood – play essential roles in wood decomposition and nutrient cycling. They are, however, strongly affected by intensive forest management making them threatened (e.g. Gossner et al. 2013). Their high diversity, well-resolved ecology, and strong association with deadwood and TreMs (since about 60 % of TreMs contain decaying wood) make them an informative model group for studying biodiversity responses to forest stand decline (Beudert et al. 2015; Bouget et al. 2019; Kozák et al. 2021; Cours et al. 2021, 2022).

To date, most studies have examined direct biodiversity responses to disturbance severity or affected vs. unaffected stands (Viljur et al. 2022). However, the indirect or cascading pathways linking disturbance legacies to community responses remain understudied. Identifying these mechanisms is essential for interpreting biodiversity patterns, refining conservation-oriented forest management, and sustain forest resilience.

Here, we assessed how saproxylic beetle taxonomic and functional α-diversity responds to stand decline across three European forest contexts: fir-dominated montane forests in the French Pyrenees, spruce-dominated montane forests in the Bavarian Alps, and oak-dominated lowland forests in the French Loire Valley. We quantified stand decline and mortality, deadwood diversity and volumes, and TreMs diversity. We hypothesized that:

(i) α-diversity and abundance of total and guild-specific saproxylic beetles would increase with deadwood and TreM quantity (i.i) and diversity (i.ii);
(ii) declining trees would primarily promote xylophagous species, whereas dead trees (snags and logs) would favor saproxylophagous species (Le Souchu et al. 2024b);
(iii) cascading pathways would differ among forest contexts due to tree species–specific disturbance legacies;
(iv) salvage logging would generate effects opposite to those observed along decline gradients.

## 2. Material and methods

### 2.1. Sampling design

We conducted our study in three different forest contexts: two in France (the Silver fir-dominated forests *Abies alba* Mill.) in French Pyrenees, hereafter in text “fir-dominated forests”, and the oak-dominated forests (mix of *Quercus petraea* (Matt) Liebl. and *Quercus robur* L.) in Loire valley, and one in Germany (the Norway spruce-dominated forests (*Picea abies* (L.) H. Karst) in Bavaria, hereafter in text “spruce-dominated forests” (Figure 1, Table 1). Along a gradient of stand decline (i.e. the proportion of dying and dead trees), we selected 56 study plots in fir-dominated forests (28 in Aure valley (Central Pyrenees; 854 to 1570 m a.s.l.; 42°51’46.8“N, 0°36’08.9“E) and 28 on Sault plateau (Eastern Pyrenees; 705 to 1557.3 m a.s.l.; 42°50’58.7“N, 2°00’41.3“E)), 15 plots in oak-dominated forests (6 in Orleans forest (107 to 174 m a.s.l.; 48°00’28“N, 2°09’39“E) and 9 in Vierzon forest (120 to 190 m a.s.l.; 47°15’53.9“N, 2°05’55.4“E)), and 28 plots in spruce-dominated forests (660 to 1352 m a.s.l.; 49°1’24.96“N 13°38’5.222“E; Figure 1). Fir and spruce forests included 12 and 9 plots respectively which have been salvage logged (Figure 1, Table 1), while the most valuable oak trees were regularly harvested during the decline process, which reduced the volume of deadwood in this context. In fir and oak forests, plots were set up in managed forests and the forests around were also predominantly managed. Plots in spruce forest were set up in the Bavarian Forest National Park, which is characterized by large areas of unmanaged disturbance areas but also by intervention sites with salvage logging in the buffer and former development zones of the National Park (see Müller et al., 2010; Thorn et al., 2016), as part of the BIOKLIM project (Bässler et al. 2009; Thorn et al. 2017). Decline in fir and oak forests were mainly the result of drought events (the 2003 drought being a major event in the occurrence of these declines), which mainly resulted in patch dynamics (Sallé et al. 2020). In spruce forests, the decline studied was the result of windstorms and bark beetle outbreaks, which mainly resulted in stand-replacing dynamics (Thorn et al. 2017). The most recent impactful event for our study was the effect of the windstorm Kyrill in 2007, from which most of the decline in our studied plots was the results, compounded by the subsequent bark beetle outbreaks.

**Figure 1.**
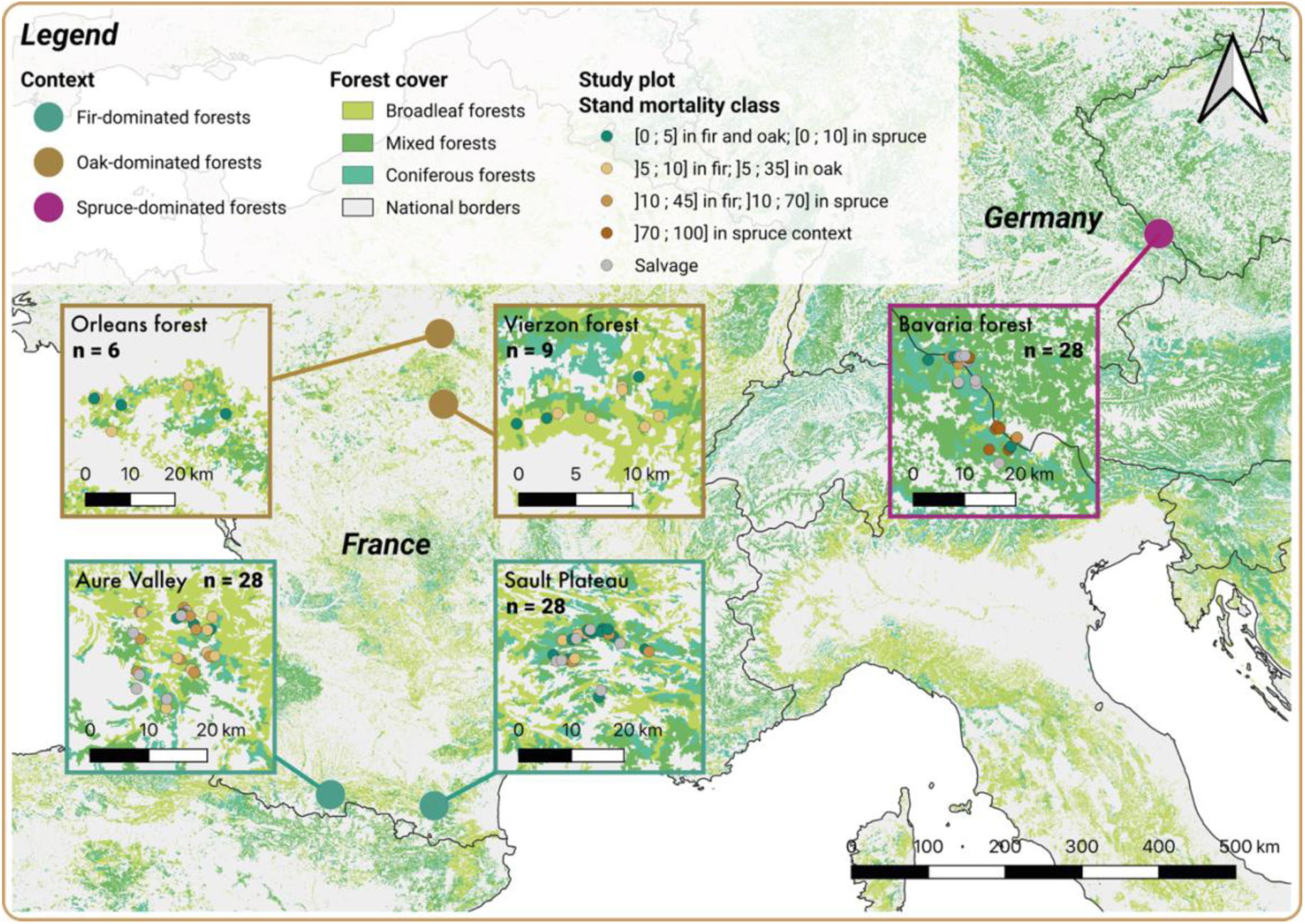
Map of the study plots and of the different study contexts.

**Table 1.** Information about the different study cases. For continuous gradients, mean, minimum, and maximum values are presented, while for salvage logging, the numbers of plots treated are presented. For more detailed information, see Figures S1 & S2.

|  | Context | Fir-dominated forests | Spruce-dominated forests | Oak-dominated forests |
| --- | --- | --- | --- | --- |
|  | Location | French Pyrenees | Bavaria (Germany) | Central France |
|  | Mean altitude | 1160 m a.s.l. | 1077 m a.s.l. | 150 m a.s.l. |
|  | Plot number | 56 | 28 | 15 |
| Environmental gradients | Stand mortality | 0.1 (0 – 0.42) | 0.55 (0 – 1) | 0.07 (0 – 0.33) |
|  | Stand decline | 0.68 (0.15 – 1) | NA | 0.33 (0 – 0.9) |
|  | Salvage logging | 12 plots | 9 plots | NA |
| Insect data set | Sampling method (traps/plot) | 2 transparent flight-interception traps | 1 transparent flight-interception traps | 2 green Lindgren traps + 1 black baited Lindgren |
|  | Saproxyllic beetles – abundance | 487 (67 – 2699) | 75 (7 – 260) | 978 (208 – 1891) |
|  | Saproxyllic beetles – Asymptotic species richness | 45 (20 – 100) | 39 (9 – 89) | 119 (49 – 186) |
| Disturbance legacies | Deadwood volume (m<sup>3</sup>.ha<sup>-1</sup>) | 50 (0 – 258) | 136 (2 – 438) | 21 (0 – 103) |
|  | Deadwood diversity | 4 (0 – 12) | 5 (1 – 9) | 1 (0 – 3) |
|  | TreM diversity | 12 (6 – 21) | 7 (1 – 15) | 8 (1 – 13) |

### 2.2. Environmental conditions

Firstly, to assess the dendrometric characteristic of each plot, we implemented a sampling protocol using relascope with 1/50 ratio (factor 1). We recorded the type (i.e. living tree, standing dead tree and lying deadwood), the tree species, the diameter at breast height (dbh) on living trees and standing dead trees higher than 4 m, and at mid-height on standing trees lower than 4 m and on lying deadwood), and the decay stage for deadwood (from 1 = fresh deadwood with full cover of bark, to 4 = soft wood with no more remaining barks). In addition, for each recorded tree, we noted the tree-related microhabitats (TreMs) on living and standing dead trees, based on the 47 types described by Larrieu et al. (2018). Mean plot area was approximately of 0.3 ha and depended on the dbh of the largest trees and their location from the center. Finally, we converted plot data into 1ha densities by allocating to every measured tree and deadwood piece, the coefficient N_d_ (Eq. 1; Pardé & Bouchon, 1988). Field data were collected in 2017 in the fir and spruce forests, and in 2020 in oak forests.

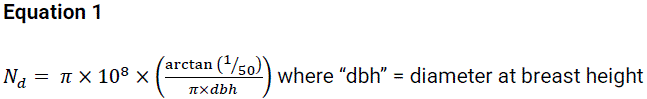

In parallel, we used eco-morphological traits related to each woody element (i.e. living trees and deadwood) and TreM, to describe environmental changes and abiotic filters. For woody elements, we directly used field measurements i.e. type of substrate (as an ordinal scale: 1 = “lying deadwood”, 2 = “standing deadwood”, 3 = “living trees”), decay stage (from 1 = “hard deadwood fully covered by bark”, to 4 = “very soft wood out of bark”), and diameter classes (see Bouget et al., 2024).

In addition, we used a list of two eco-morphological traits characterizing each TreM found in the field: (i) type of substrate bearing the TreM and (ii) saproxylic affinity. Each trait was categorized (e.g. vertical position in tree = tree base, trunk and crown) and for each TreM, we allocated a degree of occurrence in each category based on expert opinion (for additional information, see Bouget et al., 2024; data updated).

To account for the two successive processes of tree decline and death, we assessed two gradients – i.e. decline and mortality. In all forest contexts, we quantified plot-level “stand mortality” as the proportion of standing dead trees basal area relative to the total basal area of both standing dead and living trees. In parallel, we quantified the plot-level “stand decline” using ARCHI protocol (Drénou et al. 2013). This method uses tree architecture to classify tree on a decline gradient, from healthy fully functioning tree to dead tree, including three decline stages in between – i.e. resilient, stressed, and irreversible death. We applied the method on the 20 closest dominant trees from the center of each plot in fir– and oak-dominated forests. Indeed, ARCHI protocol has not been employed in spruce-dominated and therefore has not been calculated in this context (Table 1). We used the proportion of dying trees – i.e. stressed and irreversible death categories, therefore not adding dead trees – as a measure of “stand decline”.

### 2.3. Beetle sampling, identification and characterization

We sampled saproxylic beetles using different types of traps in each study context. In fir– and spruce-dominated forests, we used flight-interception traps (©Polytrap). They consisted in crossed pair of transparent plastic panes (40 × 60 cm) above a funnel conducting into a container filled with an unbaited preservative (50% propylene glycol and 50% water with a drop of detergent). There were two interception-traps in fir forests, each at least 20 m from the other, placed around the center of each study plot (Cours et al. 2022), while there was only one interception-trap in spruce forest, placed at the center of each study plot (Cours et al. 2021). The traps were suspended roughly 1.5 m above the ground and were sampled every month from mid-May to mid-September 2016 in spruce forests and 2017 in fir forests. In oak forests, we used a combination of two types of traps in each plot, two green multi-funnel traps and one black multi-funnel trap baited with volatiles (a blend of cerambycid pheromones, ethanol and alpha-pinene, hooked to the trap, see Roques et al., 2023) to sample different families of saproxylic beetles (Sallé et al. 2020). As in fir and spruce forests, collectors were filled with unbaited preservative. Each trap was installed in a separate tree at least 50 m from each other, around the plot center. These traps were suspended in the tree canopy, approximately 15 m above the ground. The traps were sampled every month from mid-May to the end of August 2020. All the captured saproxylic beetles were identified at the highest possible taxonomic level – i.e. 87% at the species level, 1% at the genus level and 12% at the family level. We retained the genera in which 100% of regional species are known to be saproxylic, and families in which at least 75% of regional species are known to be saproxylic (Bouget et al. 2019). Since we could not identify most of Staphylinidae species, and since their exclusion is not considered to have a significant impact on the analysis of the response of saproxylic beetles (Parmain et al. 2015), we excluded them from the data sets.

We characterized each saproxylic beetle species by seven functional traits: the preferred (i) deadwood diameter and (ii) stage of decay, (iii) canopy closure, (iv) the mean body size, (v) their trophic regime, (vi) the used TreM, and (vii) whether they are flower-visitors or not. The three first indices were niche traits based on species occurrence data, i.e. class of deadwood diameter and stage of decay in which species were recorded (see Gossner et al., 2013; Janssen et al., 2017). Mean body size was based on direct measurements from the FRISBEE database (Bouget et al. 2019), as well as the larval feeding guild, the larval microhabitat, and the adult floricolous behavior. For further analysis, we specifically focused on “Xylophagous”, “Saproxylophagous”, and “Zoophagous” regarding the larval feeding guild, and on “Xylofungicolous” and “Cavicolous” species regarding the TreM used by the larva (Table 2).

**Table 2.**
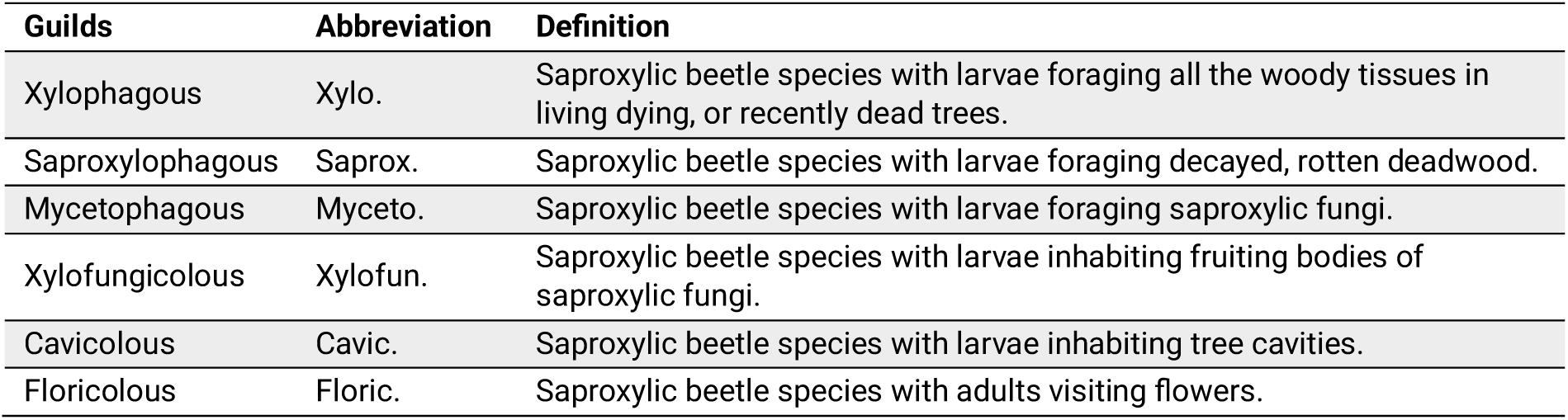
The different guilds studied in abundance and species richness, used abbreviations in figures and their definitions (Bouget et al. 2005).

### 2.4. Statistical analysis

Data analysis was conducted with R software 4.5.1 (R Core Team 2025).

Regarding the explanatory variables, we used the quantitative eco-morphological traits from woody elements and TreMs to compute community weighted means (CWM), functional dispersion (FDis) for each of them, and/or overall functional richness and dispersion, as predictors, using FD R-package (Laliberté et al., 2014; for more information, see Bouget et al., 2024). We also calculated the volume of the different types of deadwood, i.e. standing and lying, low and high decay level (i.e. level ≥ 3 on a scale from 1 to 4), small and large (dbh ≥ 40 cm), and deadwood diversity combining all these features with tree species (Siitonen et al. 2000). In addition, we calculated the total TreM diversity (based on the 47 types described by Larrieu et al., 2018), and the density per hectare of trees bearing fruiting bodies of saproxylic fungi, tree injuries and exposed sapwood, rot-holes, and saproxylic TreMs known to be important for saproxylic beetles (Stokland et al. 2012; Larrieu et al. 2018).

Regarding the response variables – i.e. related to saproxylic beetles –, we used the community matrices and the trait databases – i.e. the preference in deadwood diameter, stage of decay and canopy closure, and the mean body size – to calculate the functional indices. First, we removed species related to broadleaf tree species in both coniferous-dominated contexts (and vice versa for oak-dominated context). The functional richness (FRic), RaoQ, and evenness (FEve) were calculated using the *FD* R-package, implementing abundance community matrices (Laliberté et al. 2014). We also calculated the CWM and FDis of each of the quantitative traits used for the functional indices. In addition, we calculated the taxonomic diversity and abundance for the entire communities, but also for each ecological guilds (Bouget et al. 2019), defined by larval feeding guild (i.e. xylophagous, saproxylophagous, and mycetophagous), larval microhabitat (i.e. xylofungicolous and cavicolous), and adult behavior (i.e. floricolous, see Table 2). Finally, we used *iNEXT* R-library (Hsieh & Chao 2020) to calculate taxonomic asymptotic level of diversity (i.e. asymptotic species richness, Hill number *q* = 0), therefore disentangling from the effect of abundance. We also included the calculation of asymptotic Shannon diversity index for all species (Hill number *q* = 1). Hereafter, every index of species richness or Shannon are the result of the asymptotic calculations.

Finally, to calculate the cascading effects of stand decline and mortality, and of salvage logging on environmental conditions and saproxylic beetles, we employed piecewise Structural Equation Modelling (SEM), using the *piecewiseSEM* R-library, version 4.1.2 (Lefcheck et al. 2020). In our SEMs, hierarchical relationships were the direct effects of stand decline and mortality, and salvage logging on trees (i.e. woody elements) and subsequently on TreMs, which all may had effects on saproxylic beetles. The models implemented in the SEM were a mix of linear mixed models, for which we log-transformed the variables that required it, and generalized linear mixed models for the abundance and species richness of saproxylic beetles, using respectively the negative binomial (after testing for residual over-dispersion, using “check_overdispersion” function from *performance* R-library, Lüdecke et al., 2020) and Poisson distribution families. We also added a random variable using site – i.e. Aure and Sault in fir context, and Orléans and Vierzon in oak context; Figure 1 –, and the plot altitude as fixed covariable when significant in model to account for the within-site biogeographic context. In addition, to account for the very high correlation between the two response variables, we systematically added all species richness within beetle functional richness models. We tested each model using the function “check_model” from *performance* R-library (Lüdecke et al. 2020), to test for residual linearity, predictor collinearity, and potential outliers. We divided the analysis in two datasets: (i) a first dataset for testing both the effects of stand decline (only in fir– and oak-dominated forests) and mortality, therefore not using salvage logged plots, and (ii) a second dataset for testing the effect of salvage logging, therefore using only salvage logged plots and unsalvaged plots from the most severe mortality class in each coniferous context – i.e. 10-45% in fir-dominated forests and 70-100% in spruce-dominated forests (Figure 1). In models, salvage logging has been used as a binary variable – i.e. absence–occurrence of salvage logging. In spruce-dominated forests, since we only tested the effect of either forest mortality or salvage logging – i.e. only one tested effect in each part of analysis – on woody elements, we used the 5% α-error *p*.value to determine the significant effects. In fir– and oak-dominated forests, we used a 2.5% α-error *p*.value to test the effects of both stand decline and mortality effects as a strict adjustment of the *p*.value (0.05/2). For TreMs, we tested either forest mortality and decline, or salvage logging, effects on woody element metrics (i.e. 15 or 14 tested effects). Likewise for biodiversity, we tested for the potential effects of all explanatory variables (i.e. 25 or 24 tested effects). Therefore, we respectively adjusted the α-error *p*.value to 0.0033 or 0.0036 (as the result of 0.05/15 and 0.05/14), and 0.002 or 0.0021 (as the result of 0.05/25 and 0.05/24). In addition, we also tested for the direct effect of stand decline and mortality, and salvage logging on all the response variables (see bottom part of SEM figures), but also on the abundance of the species summing at least 10 individuals and occurring in at least 10% of the plots in each of the three contexts (Figure 5). We used the same model structuration as in the SEM for the species abundance, i.e. the tested explanatory variable, a random site factor in fir– and oak-dominated forests, and the negative binomial distribution family.

## 3. Results

We found 27,299 individuals from 127 beetle species in fir-dominated forests, 2,341 individuals from 103 species in spruce-dominated forests and 14,676 individuals from 238 species in oak-dominated forests.

### 3.1. Cascading effects of forest decline and salvage logging on taxonomic and functional α-diversity of saproxylic beetles

We found complex patterns of cascading effects of stand decline, mortality and salvage logging on saproxylic beetle communities. Those cascading effects were strongly influenced by the forest context and/or the beetle guild. Nonetheless, we observed several major effects. TreM trait dispersion was promoted by stand decline in fir-dominated forests and by stand mortality in spruce-dominated forests, enhancing overall beetle abundance in both contexts, although only as a trend in the fir-dominated forests (Figures 2 and 3). It also increased the abundances of xylophagous and mycetophagous species in fir-dominated forests (Figure 2).

**Figure 2.**
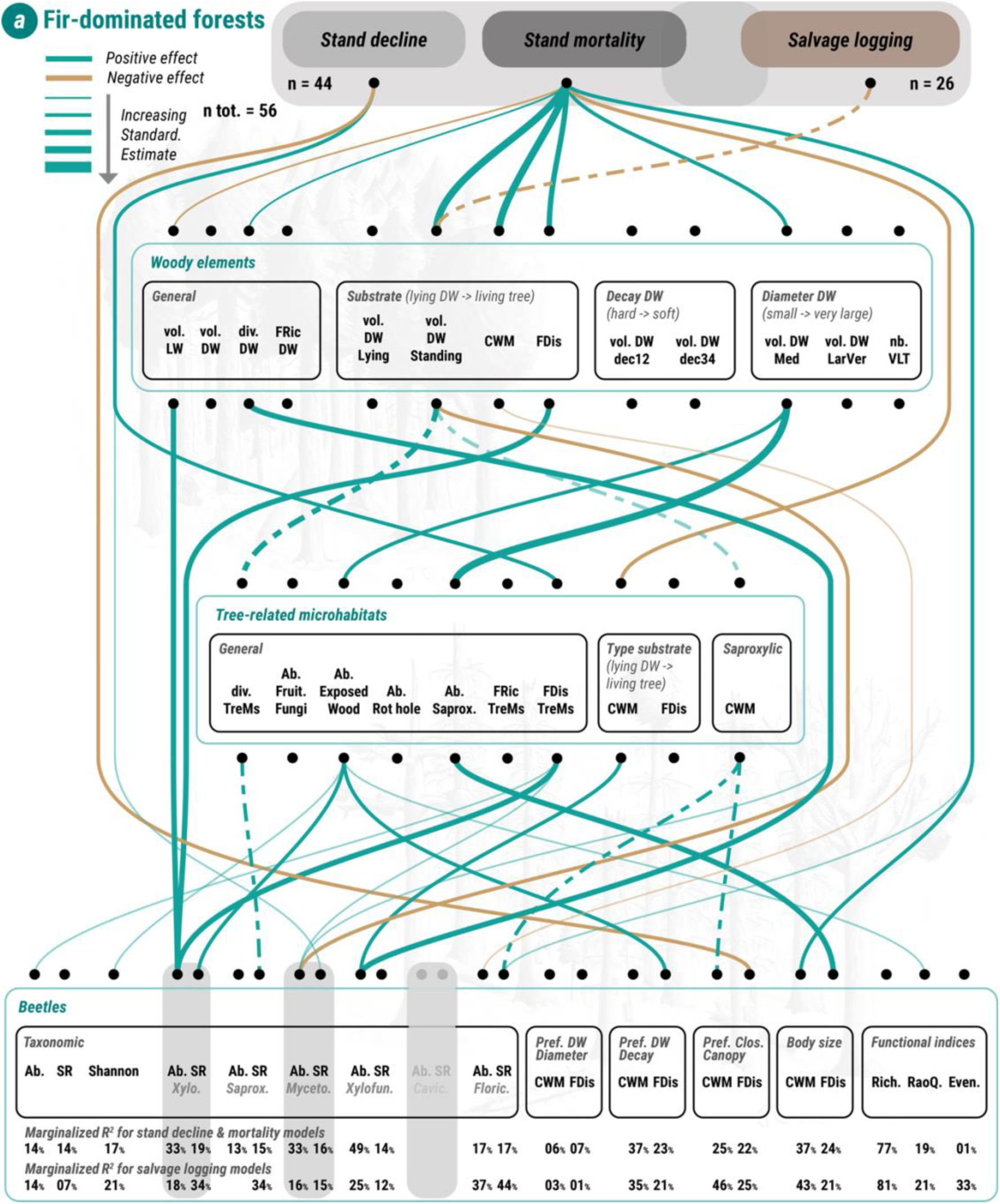
Results from structural equation modelling (SEM) for fir-dominated forests (Pyrenees region, drought disturbance). The structure of SEM is “forest decline –> woody elements –> tree-related microhabitats –> saproxylic beetles”. *P*.values are adjust based on the number of potential effects – see Methods. Effects with *p*.value superior to the adjusted *p*.value but inferior to 0.01 before correction are shown in transparent colors. Green lines show positive effects and brown lines negative ones. Solid lines show effects from stand decline & mortality models, while dashed lines show effects from salvage logging ones. Line widths illustrate the magnitude of effects (standardized estimates). The percentages are the marginalized R-squared for stand decline & mortality models (top) and salvage models (bottom). All species richness indices and the Shannon index are asymptotic levels of diversity (iNEXT calculation, see Methods). *“vol.” = “volume”; “LW” = “Living Wood”; “DW” = “Deadwood”; “Ab.” = “Abundance”; “SR” = “Species Richness”; “FDis” = “Functional Dispersion”; “FRic” = “Functional Richness”; “Rich.” = “Richness”, “Even” = “Evenness”*

**Figure 3.**
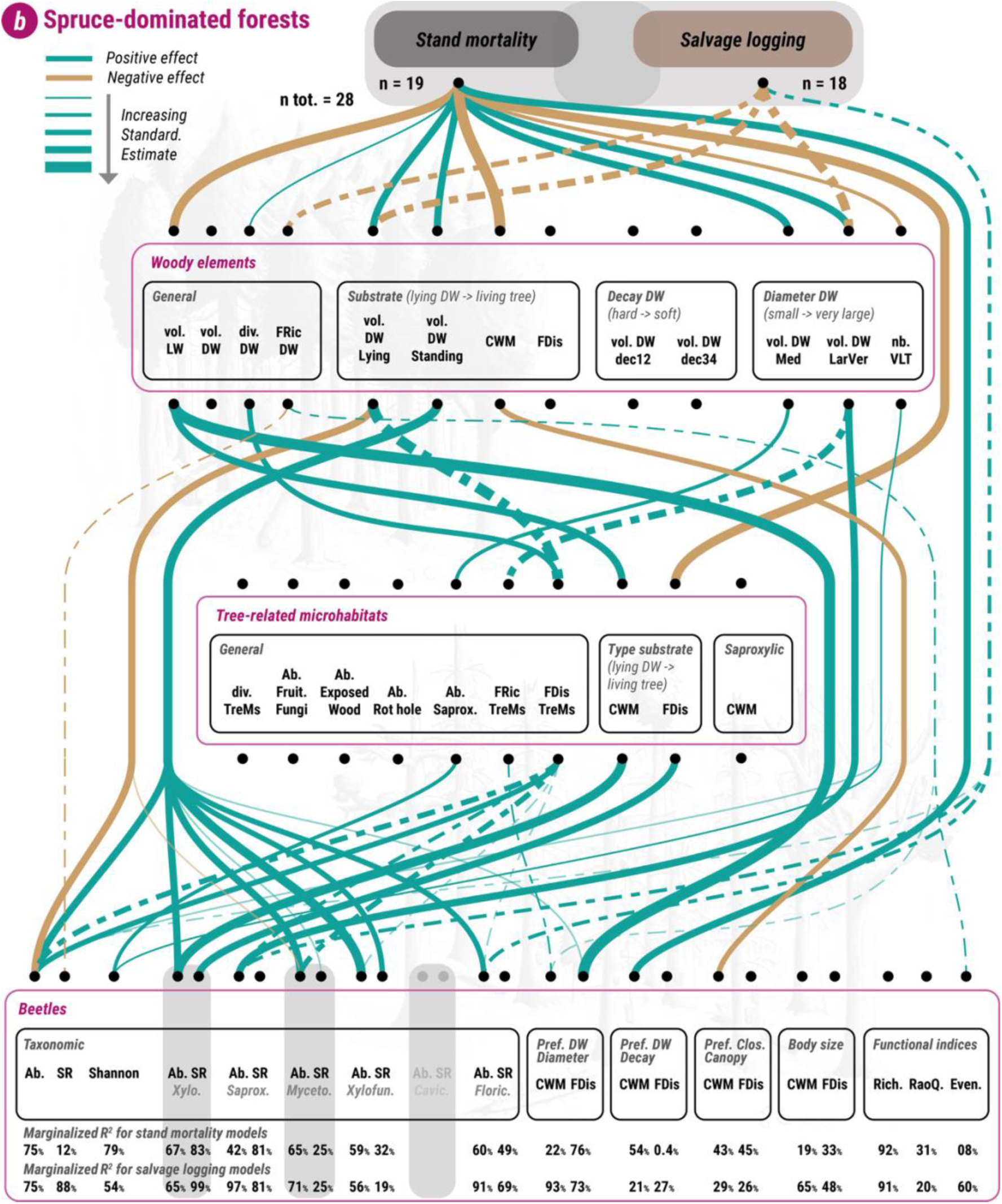
Results from structural equation modelling (SEM) for spruce-dominated forests (Bavaria region, windstorm and insect-outbreak disturbances). For additional information, see legend of Figure 2.

In addition, the density of trees bearing certain types of TreMs, also promoted by either stand decline or mortality, positively influenced certain guilds of beetles and community traits. In fir-dominated forests, the density of trees bearing exposed wood TreMs favored overall Shannon index and the species richness of xylophagous species (Figure 2). They also increased the dispersion of the beetles’ preference in deadwood decay and slightly increased beetle functional RaoQ (Figure 2). In oak-dominated forests, exposed wood TreMs density also promoted the abundance in flower-visitors (Figure 4). Moreover, density of trees bearing saproxylic TreMs increased beetle body size dispersion in fir-dominated forests (Figure 2), while it also promoted overall Shannon diversity index in spruce-dominated forests (Figure 3).

**Figure 4.**
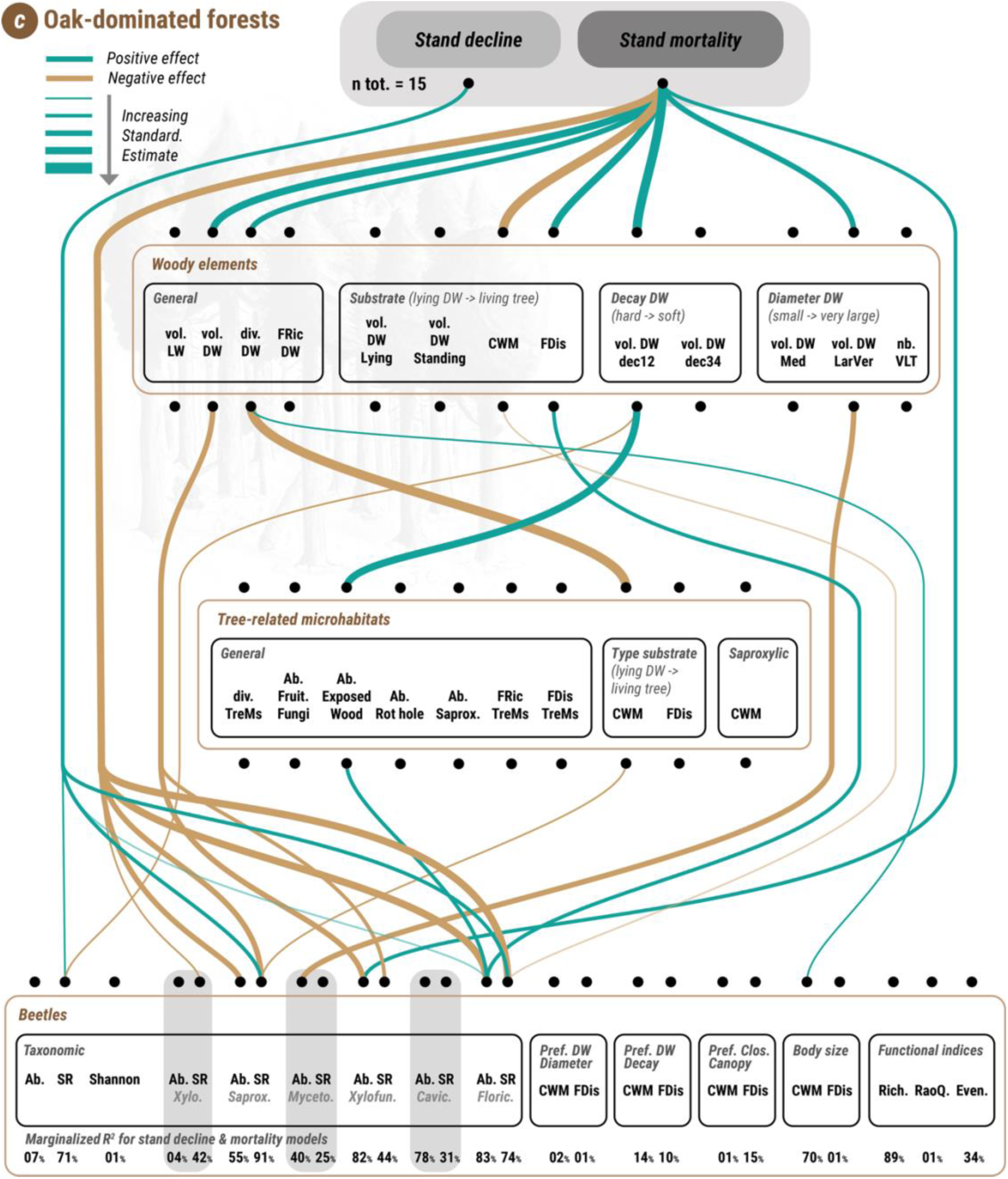
Results from structural equation modelling (SEM) for oak-dominated forests (Loire region, drought disturbance). For additional information, see legend of Figure 2.

Changes in TreM trait values also led to changes in saproxylic beetle communities. The decrease in value of substrate type bearing TreMs – i.e. less TreMs borne by living trees – had a negative effect on the abundance of xylofungicolous species in fir context (Figure 2), and xylophagous species in spruce context (Figure 3). Similarly, the reduction of the dispersion of type of substrates bearing TreMs caused by stand mortality in spruce context, led to a decrease of xylophagous species richness (Figure 3). In oak-dominated forests, contrary to spruce context, the promotion of TreMs borne by deadwood due to stand mortality, benefitted the species richness of saproxylophagous species (Figure 4).

The negative effects of salvage logging on saproxylic beetle guilds were also linked to a reduction in certain TreMs. In fir-dominated forests, the reduction in TreM diversity and saproxylic TreMs caused by standing deadwood harvest, negatively affected the richness in saproxylophagous and flower-visiting species (Figure 2). Additionally, reduction in saproxylic TreMs led to a reduction in community averaged preference for closed forest (Figure 2). In spruce-dominated forests, the marked decrease of TreM trait dispersion caused by log harvesting negatively affected overall beetle abundance, as well as the abundances in saproxylophagous and mycetophagous species, and, to a lesser extent, those of xylofungicolous and flower-visiting species (Figure 3). Additionally, in spruce context, reduction in TreM trait richness caused by salvage logging led to a reduction in community averaged preference in deadwood diameter (Figure 3).

Despite numerous cascading effects observed via TreMs, several other observed paths were directly due to changes in trees and deadwood resources. In fir-dominated forests, we found that the decrease in volume of living trees caused by stand mortality might have negatively affected the abundance in xylophagous (Figure 2). Such living tree volume reduction in spruce context led to a decrease in dispersion of the community preference in deadwood diameter (Figure 3). Otherwise, deadwood diversity promoted the abundance in xylofungicolous species, while increasing dispersion of substrate type favored directly the abundance in xylophagous species in fir context (Figure 2). In oak-dominated forests, the increasing deadwood diversity caused by stand mortality, led to an increase in averaged community body size (Figure 4). In spruce-dominated forests, while the volume of logs negatively affected the overall beetle abundance, the increased volume in standing deadwood due to stand mortality markedly benefited to the overall abundance, as well as the abundances in xylophagous, mycetophagous, xylofungicolous, and floricolous species (Figure 3). The amount of standing deadwood also benefited the richness of xylofungicolous (Figure 3). In addition, the increasing amount of large and very large deadwood promoted the abundance of saproxylophagous in spruce-dominated forests (Figure 3), while the reduction in very large trees caused by stand mortality might have slightly reduced the overall Shannon index (Figure 3). In contrast, in oak-dominated forests, the increased amount of large and very large deadwood has negatively affected the abundance in mycetophagous species (Figure 4). Furthermore, in spruce-dominated forests, the substrate changes toward fewer living trees and more deadwood have led to an averaged community preference for more closed forest environment (Figure 3). Such shift in woody substrate led to higher richness in flower-visiting species in oak-dominated forests (Figure 4). Likewise, the more diverse substrates found in high mortality stands, favored the abundance in flower visiting beetles (Figure 4). In addition, we found that overall deadwood volume negatively affected the xylofungicolous guild in oak context (Figure 4).

Finally, we also found several direct effects from stand decline and mortality, or by salvage logging occurrence. In fir-dominated forests, stand decline directly reduced the dispersion of community canopy closure preference, while it also slightly favored the richness in mycetophagous species (Figure 2). Moreover, in fir context, stand mortality directly led to an increase in averaged beetle community body size (Figure 2). In spruce-dominated forests, stand mortality led to a direct increase in averaged community preference in deadwood decay (Figure 3). We found a very contrasting pattern in oak-dominated forests, with stand decline promoting overall species richness, as well as richness of saproxylophagous and flower-visiting species, while stand mortality negatively affected xylophagous, saproxylophagous, xylofungicolous, and flower-visiting species richness (Figure 4). In addition, stand mortality in this context reduced the abundances in saproxylophagous and flower-visiting species (Figure 4). In contrast, stand mortality seems to have promoted the abundance in xylofungicolous species (Figure 4). Furthermore, salvage logging in spruce-dominated forests promoted directly the abundances in saproxylophagous and flower-visiting species (Figure 3).

### 3.2. Direct effects of forest decline and salvage logging on the most abundant saproxylic beetle species in each context

In fir-dominated forests, effects of either stand decline or mortality were generally positive on the abundances of the most abundant saproxylic species, with negative effects only found for several taxa – i.e. mainly *Epuraea* species and *Pediacus dermestoides* (Figure 5a). We also noticed that only *Nacerdes gracilis* positively responded to both stand decline and mortality (Figure 5a). In addition, we found that salvage logging had positive effects on *Pityophthorus pityographus* and *Trypodendron lineatum* (Figure 5a), and therefore, no negative effect was observed on the most abundant species.

**Figure 5.**
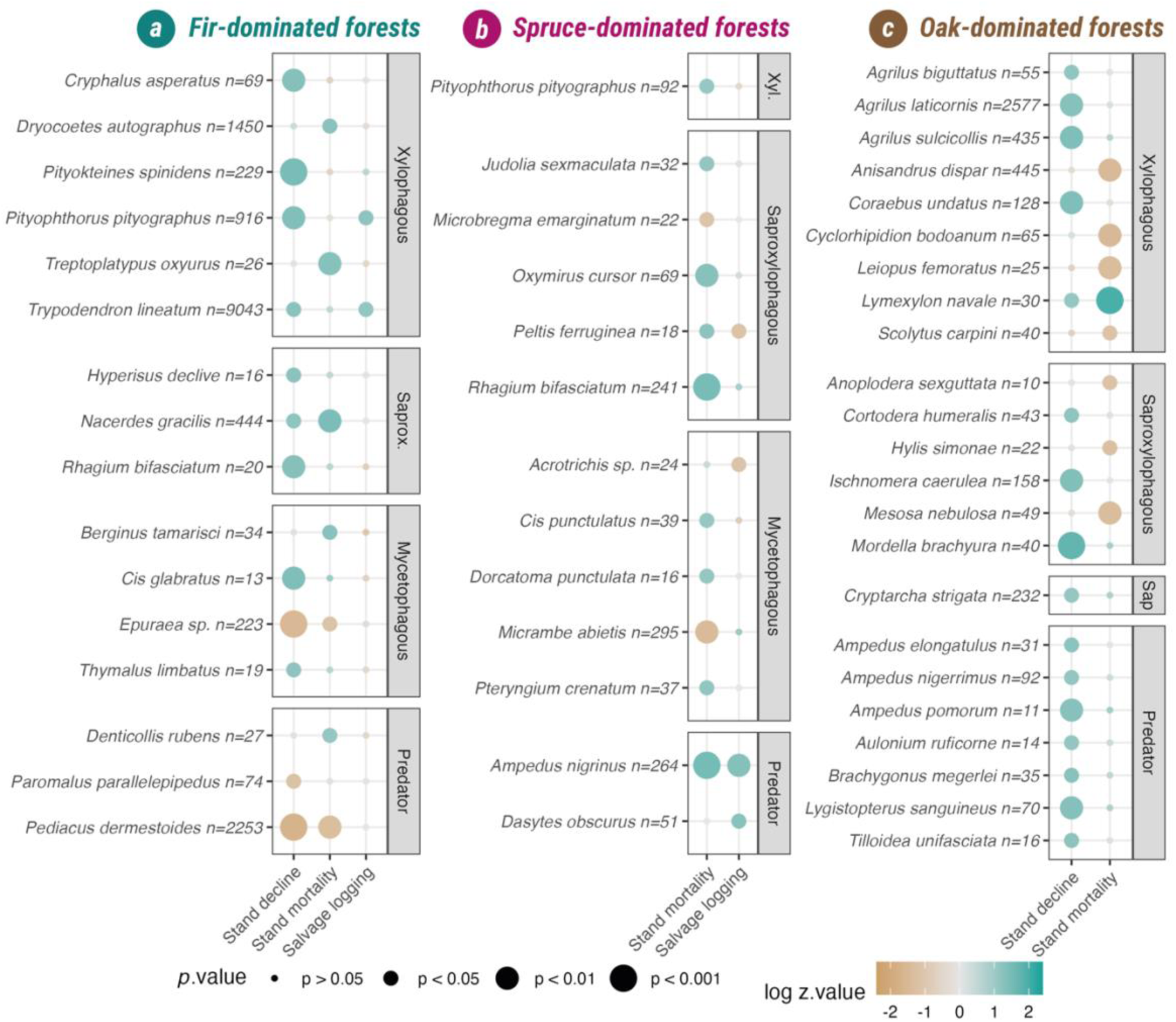
Species-scale responses (species or genus) in abundance to the mortality and decline severity, and salvage logging. Selection of taxa present in at least 10% of sites and summing at least 10 individuals in each context. Results extracted from Negative Binomial glmm including site random factor. Number of captured individuals next to taxa name. *“Xyl.” = “Xylophagous”; “Saprox.” = “Saproxylophagous”; “Sap” = “Sap-feeders”*

In spruce-dominated forests, we mostly found positive effects of forest mortality (e.g. for *Oxymirus cursor* or *Rhagium bifasciatum*), but also negative effects on *Microbregma emarginatum* and *Micrambe abietis* (Figure 5b). Salvage logging in spruce-dominated forests, promoted the abundances of *Ampedus nigrinus* and *Dasytes obscurus*, while also decreased abundances of *Peltis ferruginea* and *Acrotrichis* species (Figure 5).

In oak-dominated forests, forest decline had overall positive effect on the most abundant taxa while forest mortality has inverse effect, negatively affecting the abundance of most of the taxa, excepted for the abundance of *Lymexylon navale* (Figure 5c).

## 4. Discussion

### 4.1. Cascading effects of stand decline on beetle taxonomic and functional diversity

While some differences emerged between the effects of stand decline and stand mortality – and across the different forest contexts – both processes had a general positive influence on saproxylic beetle communities in conifer-dominated stands. This supports our first hypothesis and aligns with previous findings (Sallé et al. 2020; Cours et al. 2021, 2022; Viljur et al. 2022). However, the generally positive effect observed in coniferous contexts was more contrasted in oak-dominated forests where stand decline had a positive effect overall, whereas further stand mortality had the opposite effect (see e.g. Le Souchu et al. 2024a in the same context). Beyond confirming earlier results, our study provides new insights into the mechanisms driving community changes, identifying key cascading effects mediated by changes in both deadwood and TreM availability.

#### 4.1.1. TreMs and deadwood diversity as important and consistent resources for saproxylic beetles in context of forest stand decline

TreMs captured many of the cascading effects shaping saproxylic beetle communities during stand decline. In both coniferous-dominated forests, greater dispersion of TreM traits was associated with higher beetle abundance. Likewise, the density of trees bearing exposed wood TreMs (in fir stands) or saproxylic TreMs (in spruce stands) increased the asymptotic Shannon diversity index. In oak context, a higher density of trees bearing exposed wood promoted the abundance of floricolous species.

The relationship between habitat heterogeneity and biodiversity is well established. The *habitat-heterogeneity hypothesis* predicts that diverse habitats promote species richness through niche differentiation, whereas the *more-individuals hypothesis* suggests that increased resource availability first boosts abundance, which then secondarily enhance species richness (Clarke & Gaston 2006; Seibold et al. 2016; Müller et al. 2018). Our results rather support the more-individual mechanism: higher TreM trait dispersion likely reflects greater resource availability for saproxylic beetles (Müller et al. 2018). Similarly, the asymptotic Shannon index primarily tracked changes in dominant species, highlighting an effect on abundance rather than species richness.

Among the TreMs promoted by stand decline and/or mortality, exposed wood and injuries were the most influential (Bouget et al. 2024; Chowdhury et al. 2024; Przepiora & Ciach 2025). They increased xylophagous species richness and dispersion of beetle deadwood decay preference in fir-dominated forests, and boosted floricolous abundance in oak-dominated forests. These microhabitats likely reflect the transition between within early beetle succession, where injured dead trees host overlapping populations of species feeding on inner bark (e.g. bark beetles) or sapwood (e.g. ambrosia beetles). They may also serve as shelter for certain other groups, such as the floricolous beetles in oak forests. However, the density of exposed-wood trees may also act as a proxy for unmeasured factors.

Deadwood diversity likewise emerged as a key factor. It promoted saproxylic TreMs in fir-dominated forests, increased TreM trait dispersion in spruce-dominated forests, and was positively associated with beetle body size and indirectly with saproxylophagous richness in oak-dominated forests. Previous studies have identified deadwood diversity as a consistent predictor of saproxylic beetle diversity in temperate forests (Bouget et al. 2013). A diverse deadwood pool provides high-quality substrates for specialists (e.g. large-diameter logs; Lachat et al., 2013; Similä et al., 2003), and enhanced niche differentiation (Seibold et al. 2016). For instance, large and very large deadwood volumes supported the saproxylophagous abundance in spruce-dominated forests.

#### 4.1.2. Stand decline supports specialized saproxylic beetles and promotes threatened species

Stand mortality increased average beetle body size, directly in fir-dominated forests and indirectly via enhanced deadwood diversity in oak-dominated forests. Large-bodies saproxylic beetles are disproportionately red-listed (Seibold et al. 2015; Hagge et al. 2021), and our results therefore indicate that forest decline and mortality may benefit species that are currently declining in managed European forests (Cours et al. 2021; Busse et al. 2022).

In spruce-dominated forests, community deadwood-diameter preferences became more constrained toward larger deadwood along the stand mortality gradient (*p* = 0.04, not retained after *p*-value adjustment), consistent with findings by Cours et al. (2021). This pattern suggests increasing community specialization on large-diameter substrates as stand declines. Stand decline also promoted species associated with more advanced deadwood decay stages, directly in spruce-dominated forests and indirectly in fir-dominated forests through trees bearing exposed wood TreMs. Previous work similarly found that landscape-scale decline in fir forests favored communities relying on advanced decay stages and large deadwood (Cours et al. 2022).

These trends parallel responses observed under low-intensity forestry, where beetle communities shift toward larger and more decayed substrates (Gossner et al. 2013). Together, they highlight the role of stand decline in fostering specialized and conservation-priority saproxylic species.

#### 4.1.3. Standing deadwood as key habitat in declining spruce-dominated forests

In spruce-dominated forests, standing deadwood – snags and dead trees – emerge as a critical structural feature shaping saproxylic beetle communities. Standing deadwood increased overall species richness and enhanced the abundance and/or richness of mycetophagous, xylofungicolous, and floricolous beetles, underscoring their central role in supporting saproxylic biodiversity in spruce forests (Bouget et al. 2014).

Snags provide heterogeneous microhabitats due to variability in decay stages, humidity, sun exposure, diameter, height, and TreM diversity. This structural heterogeneity supports diverse resources promoting both beetle abundance and species richness (Bouget et al. 2012, 2014; Parisi et al. 2018; Kozák et al. 2021; Cours et al. 2021). Their importance is particularly pronounced in coniferous forests, where snags are often the primary source of cracks and polypores – microhabitats that are rare on living trees (Larrieu & Cabanettes 2012). For instance, *Cis punctulatus*, a mycetophagous species preferring tall snags (Hjältén et al. 2010), responded positively to spruce stand mortality. In fir-dominated forests, although the negative relationship between snag volume and mycetophagous beetle abundance is difficult to interpret, our results indicate that snags enhance TreM diversity under salvage logging, thereby supporting saproxylophagous richness.

In contrast, lying deadwood exhibited opposite effects in spruce-dominated forests. Despite similar decay stages and diameters to standing deadwood (Figure S7), greater log volumes were associated with lower overall species richness, particularly among xylofungicolous species. This finding is unexpected given that logs typically complement snags in enriching local species pool (Jonsell & Weslien 2003; Hjältén et al. 2010; Bouget et al. 2012; Graf et al. 2022). Because snag and log volumes were not negatively correlated (Figure S3c), reduced richness associated with logs likely reflect unmeasured ecological processes, such as resource specialization or species interactions. Future work should assess species-level preferences for snags versus logs and investigate potential negative interactions – e.g. competition or habitat exclusion – that may structure communities in declining spruce stands.

### 4.2. Consistencies and contrasts between the effects of canopy decline and tree mortality

Our results support the hypothesis that stand decline and stand mortality exert distinct influences on saproxylic beetle communities. In fir-dominated forests, stand decline initiated cascading effects through increased TreM trait dispersion, which boosted overall beetle abundance and particularly favored xylophagous species. In oak-dominated forests, although overall responses of xylophagous community were not significant, higher proportions of declining trees promoted several early-stage xylophagous beetles (i.e. *Agrilus biguttatus*, *A. laticornis*, or *A. sulcicollis*, and *Coraebus undatus*; Sallé et al., 2020; Le Souchu et al. 2024b). These patterns indicate that stand decline represents an early succession stage in which stressed but still-living trees offer highly suitable substrates for opportunistic colonizers (Stokland et al. 2012; Thorn et al. 2016; Wermelinger et al. 2025), particularly in drought-affected systems where tree death unfolds slowly.

In contrast, stand mortality generated divergent cascading effects depending on forest context. In coniferous-dominated forests, mortality advanced successional dynamic by promoting saproxylophagous species (e.g. *Nacerdes gracilis*, *Rhagium bifasciatum*) that typically colonize after xylophagous pioneers. In oak-dominated forests, however, mortality produced opposite outcomes relative to stand decline: it reduced overall species richness – including saproxylophagous richness – and decreased the abundance of mycetophagous and xylofungicolous species. Several dominant saproxylophagous species (e.g. *Mesosa nebulosa*) also responded negatively to oak mortality.

These contrasting patterns may stem from inherent differences in wood physical and chemical properties. Oak decomposition is slower and shaped by distinct wood-decay fungal pathways (mostly fibrous white-rot) compared with conifers, which typically mostly host cubic brown-rot (Kahl et al. 2017). Although short-term effects of fungal communities on beetles appear limited (Sbaraglia et al. 2025), they could exert long-term effects via priority effects (Weslien et al. 2011). Microclimatic context likely further amplifies these contrasts: in coniferous forests at high altitudes and/or latitudes, canopy openings may warm deadwood while abundant precipitations maintains wood moisture—conditions favorable for colonization (Seibold et al. 2016; Harmon et al. 2020; Sbaraglia et al. 2025). Conversely, in warm, dry lowland oak forests, openings likely accelerate deadwood desiccation, reducing suitability for many saproxylic taxa (Harmon et al. 2020). Finally, ongoing logging in oak forests may negatively affect saproxylic beetles and confound results (Thorn et al. 2018).

### 4.3. Contrasted conditions and cascading effects between forest contexts

Stand mortality was most severe in spruce-dominated stands, which showed the highest deadwood volumes—up to 438 m^3^.ha^−1^ (Figure S1)—and the sharpest contrast between low and high decline levels (see Table 1). In contrast, TreMs diversity was greatest in fir-dominated forests (Figure S1). Although oak-dominated forests displayed high TreM trait diversity, mean TreMs abundances were consistently lower than in coniferous contexts (Figure S1), despite oak’s general tendency to host more and richer TreMs (Courbaud et al. 2022). This discrepancy likely reflects the effects of ongoing salvage logging in oak context. These structural contrasts generated distinct cascading effects on saproxylic beetle communities across forest contexts (Sallé et al. 2021; Cours et al. 2021, 2023; Viljur et al. 2022).

Each declining context also supported distinct beetle communities aligned with different successional stages. In spruce-dominated forests, the rapid and severe mortality triggered by *Ips typographus* occurred several years before sampling. Consequently, we observed only one dominant xylophagous species – *Pityophthorus pityographus* – and a negative effect of mortality on xylophagous richness, likely due to the loss of living substrates bearing TreMs. This pattern parallels observations from Switzerland, where xylophagous beetles peak in the five years following windstorm and subsequently collapse to near-zero levels (Wermelinger et al. 2025). In contrast, drought-induced decline in fir– and oak-dominated forests progressed slowly, with trees dying over several years. This gradual trajectory allows temporal overlap between early– (e.g. xylophagous) and late-successional (e.g. saproxylophagous) beetle groups, generating more continuous community transitions.

Flower-visiting beetles also showed strong context dependency. Their relative abundance was lowest in fir-dominated forests than in the two other contexts, where they increased with decline (Figures 3 & 4). These differences likely reflect variations in canopy openness. The slow drought-driven decline in fir-dominated forests produced more limited canopy opening compared with the more severe decline in spruce forests, and the more actively logged oak forests (Cours et al. 2021). Canopy openness is a major driver of beetle assemblages (Bouget et al. 2013), and forest insect communities more broadly (Sire et al. 2022), as sun-exposed deadwood typically hosts more species than shaded substrates (Seibold et al. 2016; Lettenmaier et al. 2022).

In spruce-dominated forests, however, tree mortality unexpectedly favored species preferring shadier environments. Although canopy loss should increase solar exposure, the dense tree regeneration might have re-created shaded conditions that benefited shade-associated beetles (Sbaraglia et al. 2025).

Overall, our results underscore the strong context dependency of stand decline and mortality effects (Catford et al. 2022). Cascading ecological responses varied depending on dominating tree species, biogeographical settings, and the natural disturbance type (Cours et al. 2023). Methodological differences – most notably, distinct trapping protocols between oak and conifer contexts – likely contributed to some observed contrasts. Nevertheless, our study clarifies how interacting structural and ecological factors drive divergent saproxylic beetle responses across European forest contexts. Given the rising frequency, severity, and extent of natural disturbances and forest stand decline globally (Allen et al. 2015; Seidl et al. 2017; Samaniego et al. 2018; McDowell et al. 2020; Viljur et al. 2022), addressing this complexity is increasingly crucial for conserving biodiversity in rapidly changing forest landscapes.

### 4.4. Contrasting effects of salvage logging on TreMs-dependant and floricolous saproxylic guilds

Salvage logging primarily affected beetle communities through reductions in TreM diversity and trait dispersion, particularly depleting saproxylic TreMs. This loss of key microhabitats and resources led to declines in overall beetle abundance, notably among saproxylophagous and mycetophagous species in spruce-dominated forests, and in saproxylophagous and floricolous species richness in fir-dominated forests. Combined with the patterns observed along decline gradients, these findings further emphasize the central role of TreMs in sustaining saproxylic beetle communities, both under conditions of habitat enrichment and habitat loss. Salvage logging specifically impacts saproxylophagous beetles because it removes decayed deadwood – a critical resource that is harvested before it can form (Priewasser et al. 2013). Our results therefore reinforce earlier evidence of the detrimental effects of salvage logging on saproxylic biodiversity via the resource depletion (Lindenmayer et al. 2008; Thorn et al. 2018).

However, salvage logging also promoted the abundances of floricolous and saproxylophagous species in spruce-dominated forests. Because many adult saproxylophagous beetles are also floricolous (e.g. numerous *Cerambycidae* species; see Figure S5), this positive effect is likely driven by their floricolous trait – i.e. saproxylophagous trophic trait is related to larval stage. This pattern agrees with previous studies showing that salvage logging promotes floricolous species (Thorn et al. 2016; Heil & Burkle 2018; Wermelinger et al. 2025). In montane forests, salvage logging creates canopy openings and soil disturbances that promote the establishment of flowering plants such as *Epilobium angustifolium*, as well as species from the *Apiaceae*, *Asteraceae,* or *Campanulaceae* families, thereby enhancing pollinator resources (Swanson et al. 2011; Thorn et al. 2016; Orczewska et al. 2019; Wermelinger et al. 2025). Nonetheless, in fir-dominated forests, we observed the contrasting decline in floricolous species richness, driven by the reduction in saproxylic TreMs.

### 4.5. Conclusion

Our study shows that forest stand decline generally enhances saproxylic beetles in European coniferous forests and identifies the cascading habitat-mediated mechanisms driving these responses. Increased heterogeneity of tree-related microhabitats generally boosted overall beetle abundance, and standing dead trees played a key role in spruce-dominated forests affected by windstorm and bark beetle outbreaks. We also demonstrated that stand decline and mortality generate distinct community responses, with different successional guilds benefiting from each. These mechanisms were strongly context dependent, underscoring the need to better understand how decline processes vary across forest contexts under a rapidly changing climate.

Declining stand islands can provide valuable conservation opportunities due to their positive effects on saproxylic beetle communities. Conversely, the harvest of habitat trees bearing diverse TreMs during salvage logging reduced resources for several beetle guilds – most notably saproxylophagous species that depend on decayed deadwood. These findings are consistent with earlier evidences of the detrimental impacts of salvage logging on forest biodiversity (Lindenmayer et al. 2008; Thorn et al. 2018, 2020).

We therefore advocate for the conservation of declining forest stand patches as temporary biodiversity refugia. At minimum, retaining TreM-rich large dead trees and logs during salvage logging operations can provide essential “lifeboats” for saproxylic beetles and help sustain biodiversity.

## Acknowledgements

This research is part of the international project CLIMTREE “Ecological and Socioeconomic Impacts of Climate-Induced Tree Dieback in Highland Forests” within the Belmont Forum Call: “Mountains as Sentinels of Change”, coordinated by C. Lopez-Vaamonde. The French team was funded by the French National Research Agency (ANR) (ANR-15-MASC-002-01). The German team was funded by the Deutsche Forschungsgemeinschaft (DFG) (MA 7249/1-1). We are thankful to Carl Moliard and Benoît Nusillard from INRAE EFNO; Wilfried Heintz, Laurent Burnel, Jérôme Molina and Jérôme Willm from INRAE DYNAFOR and Grégory Sajdak from CNPF-IDF for their field work. We are also grateful to Sylvie Ladet for her assistance in preparing the protocol and mapping work. We also thank all of those who helped us in the field: Dominique Micaux, Pierre Caillieux, Vincent Gherra, X. Fraces, Serge Alric, Xavier de Muyser, Francis Dueso, Serge Rumeau, Jean-Marie Quilès, Gilles Vergèz and David Veneau from the ONF; Jérôme Moret, Emmanuel Rouyer, Benoît Lecomte and Jean-Christophe Chabalier from the CRPF Occitanie; forest private owners Jean-Claude Marquis, Gilles Lefrançois, Marc Mesplié, Patrick Ferran, Jean-Baptiste Régné and Gilles Verdier; and Denis Sabadie and Vincent Sabadie on the Sault Plateau.

This work was also supported by Région Centre-Val de Loire Project no. 2018-00124136 (CANOPEE) coordinated by A. Sallé. We thank C. Moliard (INRAE), O. Denux (INRAE) and X. Pineau (University of Orléans) for their technical assistance. We are grateful to O. Rose (Ciidae), T. Noblecourt (Scolytinae), F. Soldati (Tenebrionidae, Carabidae), T. Barnouin (Elateridae (pars), Ptinidae (pars)), O. Courtin (Scraptiidae, Mordellidae), Benedikt Feldmann (Staphylinidae), Gianfranco Liberti (Dasytidae) and Y. Gomy (Histeridae) for their help with species identification. We are also grateful to the National Forestry Office (Office National des Forêts) and the Forest Health Service (Département de la Santé des Forêts).

JC was also partly supported by the Kone foundation (application 202105759).

## References

1. Allen CD, Breshears DD, McDowell NG. 2015. On underestimation of global vulnerability to tree mortality and forest die-off from hotter drought in the Anthropocene. Ecosphere 6:art129.

2. Bässler C, Förster B, Moning C, Müller J. 2009. The BIOKLIM Project: Biodiversity Research between Climate Change and Wilding in a temperate montane forest – The conceptual framework. Waldökologie, Landschaftsforschung und Naturschutz 7:21–33.

3. Batllori E et al. 2020. Forest and woodland replacement patterns following drought-related mortality. Proceedings of the National Academy of Sciences 117:29720–29729. National Academy of Sciences.

4. Beudert B, Bässler C, Thorn S, Noss R, Schröder B, Dieffenbach-Fries H, Foullois N, Müller J. 2015. Bark Beetles Increase Biodiversity While Maintaining Drinking Water Quality. Conservation Letters 8:272–281.

5. Bouget C, Brustel H, Nageleisen L-M. 2005. Nomenclature of wood-inhabiting groups in forest entomology: synthesis and semantic adjustments. Comptes Rendus Biologies 328:936–948.

6. Bouget C, Brustel H, Noblecourt T, Zagatti P. 2019. Les Coléoptères saproxyliques de France: Catalogue écologique illustréEditions du MNHN. Paris.

7. Bouget C, Cours J. 2025. Effects of forest dieback on deadwood patterns: Large scale trends from a cross-analysis of European databases. Journal of Environmental Management 375:124315.

8. Bouget C, Cours J, Larrieu L, Parmain G, Müller J, Speckens V, Sallé A. 2024. Trait-Based Response of Deadwood and Tree-Related Microhabitats to Decline in Temperate Lowland and Montane Forests. Ecosystems 27:90–105.

9. Bouget C, Larrieu L, Brin A. 2014. Key features for saproxylic beetle diversity derived from rapid habitat assessment in temperate forests. Ecological Indicators 36:656–664.

10. Bouget C, Larrieu L, Nusillard B, Parmain G. 2013. In search of the best local habitat drivers for saproxylic beetle diversity in temperate deciduous forests. Biodiversity and Conservation 22:2111–2130.

11. Bouget C, Nusillard B, Pineau X, Ricou C. 2012. Effect of deadwood position on saproxylic beetles in temperate forests and conservation interest of oak snags. Insect Conservation and Diversity 5:264–278.

12. Busse A et al. 2022. Forest dieback in a protected area triggers the return of the primeval forest specialist Peltis grossa (Coleoptera, Trogossitidae). Conservation Science and Practice 4:e612.

13. Catford JA, Wilson JRU, Pyšek P, Hulme PE, Duncan RP. 2022. Addressing context dependence in ecology. Trends in Ecology & Evolution 37:158–170. Elsevier.

14. Chowdhury FI, Lloret F, Jaime L, Margalef-Marrase J, Espelta JM. 2024. Deadwood and Tree-related Microhabitat’s abundance and diversity are determined by the interplay of drought-induced die-off and local climate. Forest Ecology and Management 563:121989.

15. Clarke A, Gaston KJ. 2006. Climate, energy and diversity. Proceedings of the Royal Society B: Biological Sciences 273:2257–2266. Royal Society.

16. Courbaud B et al. 2022. Factors influencing the rate of formation of tree-related microhabitats and implications for biodiversity conservation and forest management. Journal of Applied Ecology 59:492–503.

17. Cours J, Bouget C, Barsoum N, Horák J, Le Souchu E, Leverkus AB, Pincebourde S, Thorn S, Sallé A. 2023. Surviving in Changing Forests: Abiotic Disturbance Legacy Effects on Arthropod Communities of Temperate Forests. Current Forestry ReportsDOI: 10.1007/s40725-023-00187-0. Available from 10.1007/s40725-023-00187-0 (accessed May 8, 2023).

18. Cours J, Larrieu L, Lopez-Vaamonde C, Müller J, Parmain G, Thorn S, Bouget C. 2021. Contrasting responses of habitat conditions and insect biodiversity to pest– or climate-induced dieback in coniferous mountain forests. Forest Ecology and Management 482. Available from http://www.sciencedirect.com/science/article/pii/S0378112720315802 (accessed December 15, 2020).

19. Cours J, Sire L, Ladet S, Martin H, Parmain G, Larrieu L, Moliard C, Lopez-Vaamonde C, Bouget C. 2022. Drought-induced forest dieback increases taxonomic, functional, and phylogenetic diversity of saproxylic beetles at both local and landscape scales. Landscape EcologyDOI: 10.1007/s10980-022-01453-5. Available from 10.1007/s10980-022-01453-5 (accessed June 21, 2022).

20. Drénou C, Giraud F, Gravier H, Sabatier S, Caraglio Y. 2013. Le diagnostic architectural: un outil d’évaluation des sapinières dépérissantes. Forêt méditerranéenne 34:87–98. Association Forêt Méditerranéenne, 14 rue Louis Astouin, 13002 MARSEILLE ….

21. Franklin JF, Lindenmayer D, MacMahon JA, McKee A, Magnuson J, Perry DA, Waide R, Foster D. 2000. Threads of Continuity. Conservation in Practice 1:8–17.

22. Georgiev KB, Bässler C, Feldhaar H, Heibl C, Karasch P, Müller J, Perlik M, Weiss I, Thorn S. 2022. Windthrow and salvage logging alter β-diversity of multiple species groups in a mountain spruce forest. Forest Ecology and Management 520:120401.

23. Gossner MM, Lachat T, Brunet J, Isacsson G, Bouget C, Brustel H, Brandl R, Weisser WW, Müller J. 2013. Current Near-to-Nature Forest Management Effects on Functional Trait Composition of Saproxylic Beetles in Beech Forests. Conservation Biology 27:605–614.

24. Graf M, Lettenmaier L, Müller J, Hagge J. 2022. Saproxylic beetles trace deadwood and differentiate between deadwood niches before their arrival on potential hosts. Insect Conservation and Diversity 15:48–60.

25. Hagge J et al. 2021. What does a threatened saproxylic beetle look like? Modelling extinction risk using a new morphological trait database. Journal of Animal Ecology 90:1934–1947.

26. Harmon ME, Fasth BG, Yatskov M, Kastendick D, Rock J, Woodall CW. 2020. Release of coarse woody detritus-related carbon: a synthesis across forest biomes. Carbon Balance and Management 15:1.

27. Harvey JA et al. 2023. Scientists’ warning on climate change and insects. Ecological Monographs 93:e1553.

28. Heil LJ, Burkle LA. 2018. Recent post-wildfire salvage logging benefits local and landscape floral and bee communities. Forest Ecology and Management 424:267–275.

29. Hjältén J, Stenbacka F, Andersson J. 2010. Saproxylic beetle assemblages on low stumps, high stumps and logs: Implications for environmental effects of stump harvesting. Forest Ecology and Management 260:1149–1155.

30. Hsieh TC, Chao KHM and A. 2020, January 28. iNEXT: Interpolation and Extrapolation for Species Diversity. Available from https://CRAN.R-project.org/package=iNEXT (accessed November 18, 2020).

31. Janssen P, Fuhr M, Cateau E, Nusillard B, Bouget C. 2017. Forest continuity acts congruently with stand maturity in structuring the functional composition of saproxylic beetles. Biological Conservation 205:1–10.

32. Johnstone JF et al. 2016. Changing disturbance regimes, ecological memory, and forest resilience. Frontiers in Ecology and the Environment 14:369–378.

33. Jonsell M, Weslien J. 2003. Felled or standing retained wood—it makes a difference for saproxylic beetles. Forest Ecology and Management 175:425–435.

34. Kahl T et al. 2017. Wood decay rates of 13 temperate tree species in relation to wood properties, enzyme activities and organismic diversities. Forest Ecology and Management 391:86–95.

35. Kozák D et al. 2021. Historical Disturbances Determine Current Taxonomic, Functional and Phylogenetic Diversity of Saproxylic Beetle Communities in Temperate Primary Forests. Ecosystems 24:37–55.

36. Lachat T, Bouget C, Bütler R, Müller J. 2013. Deadwood: quantitative and qualitative requirements for the conservation of saproxylic biodiversity. Page 92 Integrative approaches as an opportunity for the conservation of forest biodiversity. European Forest Institute.

37. Laliberté E, Legendre P, Shipley B. 2014, August 19. FD: Measuring functional diversity (FD) from multiple traits, and other tools for functional ecology. Available from https://CRAN.R-project.org/package=FD (accessed May 8, 2020).

38. Larrieu L, Cabanettes A. 2012. Species, live status, and diameter are important tree features for diversity and abundance of tree microhabitats in subnatural montane beech–fir forests1This article is one of a selection of papers from the International Symposium on Dynamics and Ecological Services of Deadwood in Forest Ecosystems. Canadian Journal of Forest Research 42:1433–1445. NRC Research Press.

39. Larrieu L, Paillet Y, Winter S, Bütler R, Kraus D, Krumm F, Lachat T, Michel AK, Regnery B, Vandekerkhove K. 2018. Tree related microhabitats in temperate and Mediterranean European forests: A hierarchical typology for inventory standardization. Ecological Indicators 84:194–207.

40. Le Souchu E et al. 2024a. Responses of the hyper-diverse community of canopy-dwelling Hymenoptera to oak decline. Insect Conservation and Diversity DOI: 10.1111/icad.12708. Available from https://onlinelibrary.wiley.com/doi/abs/10.1111/icad.12708 (accessed January 22, 2024).

41. Le Souchu E, Bouget C, Sallé A. 2024b. Environmental drivers of local and temporal variations in the community of oak-associated borers (Coleoptera: Buprestidae). European Journal of Forest Research 143:603–616.

42. Lefcheck JS, Byrnes J, Grace J. 2020, December 9. piecewiseSEM: Piecewise Structural Equation Modeling. Available from https://CRAN.R-project.org/package=piecewiseSEM (accessed August 26, 2022).

43. Lettenmaier L, Seibold S, Bässler C, Brandl R, Gruppe A, Müller J, Hagge J. 2022. Beetle diversity is higher in sunny forests due to higher microclimatic heterogeneity in deadwood. Oecologia 198:825–834.

44. Lindenmayer DB, Burton PJ, Franklin JF. 2008. Salvage logging and its ecological consequences. Island Press. Available from https://agris.fao.org/agris-search/search.do?recordID=US201300128956 (accessed July 2, 2020).

45. Lüdecke D, Makowski D, Waggoner P, Patil I. 2020, May 3. performance: Assessment of Regression Models Performance. Available from https://CRAN.R-project.org/package=performance (accessed May 12, 2020).

46. McDowell NG et al. 2020. Pervasive shifts in forest dynamics in a changing world. Science 368:eaaz9463.

47. Müller J et al. 2018. LiDAR-derived canopy structure supports the more-individuals hypothesis for arthropod diversity in temperate forests. Oikos 127:814–824.

48. Müller J, Noss RF, Bussler H, Brandl R. 2010. Learning from a “benign neglect strategy” in a national park: Response of saproxylic beetles to dead wood accumulation. Biological Conservation 143:2559–2569.

49. Orczewska A, Czortek P, Jaroszewicz B. 2019. The impact of salvage logging on herb layer species composition and plant community recovery in Białowieża Forest. Biodiversity and Conservation 28:3407–3428.

50. Pardé J, Bouchon J. 1988. Dendrométrie2 ed. Ecole Nationale du génie rural, des Eaux et Forêts, Nancy.

51. Parisi F, Pioli S, Lombardi F, Fravolini G, Marchetti M, Tognetti R. 2018. Linking deadwood traits with saproxylic invertebrates and fungi in European forests – a review. iForest – Biogeosciences and Forestry 11:423. SISEF – Italian Society of Silviculture and Forest Ecology.

52. Parmain G, Bouget C, Müller J, Horak J, Gossner MM, Lachat T, Isacsson G. 2015. Can rove beetles (Staphylinidae) be excluded in studies focusing on saproxylic beetles in central European beech forests? Bulletin of Entomological Research 105:101–109.

53. Pickett STA, White PS. 1985. The Ecology of Natural Disturbance and Patch Dynamics. Academic Press. Available from https://linkinghub.elsevier.com/retrieve/pii/C20090029523 (accessed December 1, 2020).

54. Priewasser K, Brang P, Bachofen H, Bugmann H, Wohlgemuth T. 2013. Impacts of salvage-logging on the status of deadwood after windthrow in Swiss forests. European Journal of Forest Research 132:231–240.

55. Przepiora F, Ciach M. 2025, April 17. Bark beetles as ecosystem engineers: triggered tree mortality rearranges the assemblage of Tree-related Microhabitats in old-growth coniferous forest. bioRxiv. Available from https://www.biorxiv.org/content/10.1101/2025.04.11.648382v1 (accessed April 19, 2025).

56. R Core Team. 2025. R: A Language and Environment for Statistical Computing. R Foundation for Statistical Computing, Vienna, Austria. Available from https://www.R-project.org/.

57. Roques A et al. 2023. Worldwide tests of generic attractants, a promising tool for early detection of non-native cerambycid species. NeoBiota 84:169–209. Pensoft Publishers.

58. Sallé A, Cours J, Le Souchu E, Lopez-Vaamonde C, Pincebourde S, Bouget C. 2021. Climate Change Alters Temperate Forest Canopies and Indirectly Reshapes Arthropod Communities. Frontiers in Forests and Global Change 4:120.

59. Sallé A, Parmain G, Nusillard B, Pineau X, Brousse R, Fontaine-Guenel T, Ledet R, Vincent-Barbaroux C, Bouget C. 2020. Forest decline differentially affects trophic guilds of canopy-dwelling beetles. Annals of Forest Science 77:86.

60. Samaniego L, Thober S, Kumar R, Wanders N, Rakovec O, Pan M, Zink M, Sheffield J, Wood EF, Marx A. 2018. Anthropogenic warming exacerbates European soil moisture droughts. Nature Climate Change 8:421.

61. Sbaraglia C, Thorn S, Ambrožová L, Bässler C, Čížek L, Kozel P, Procházka J, Drag L. 2025. Sun exposure as a main driver influencing the arriving communities of saproxylic beetles on deadwood. Ecological Entomology 50:565–576.

62. Seibold S et al. 2019. Arthropod decline in grasslands and forests is associated with landscape-level drivers. Nature 574:671–674. Nature Publishing Group.

63. Seibold S, Bässler C, Brandl R, Büche B, Szallies A, Thorn S, Ulyshen MD, Müller J. 2016. Microclimate and habitat heterogeneity as the major drivers of beetle diversity in dead wood. Journal of Applied Ecology 53:934–943.

64. Seibold S, Brandl R, Buse J, Hothorn T, Schmidl J, Thorn S, Müller J. 2015. Association of extinction risk of saproxylic beetles with ecological degradation of forests in Europe. Conservation Biology 29:382–390.

65. Seidl R et al. 2017. Forest disturbances under climate change. Nature Climate Change 7:395–402. Nature Publishing Group.

66. Senf C, Buras A, Zang CS, Rammig A, Seidl R. 2020. Excess forest mortality is consistently linked to drought across Europe. Nature Communications 11:6200. Nature Publishing Group.

67. Senf C, Pflugmacher D, Zhiqiang Y, Sebald J, Knorn J, Neumann M, Hostert P, Seidl R. 2018. Canopy mortality has doubled in Europe’s temperate forests over the last three decades. Nature Communications 9:4978. Nature Publishing Group.

68. Senf C, Seidl R. 2021. Persistent impacts of the 2018 drought on forest disturbance regimes in Europe. Biogeosciences 18:5223–5230. Copernicus GmbH.

69. Siitonen J, Martikainen P, Punttila P, Rauh J. 2000. Coarse woody debris and stand characteristics in mature managed and old-growth boreal mesic forests in southern Finland. Forest Ecology and Management 128:211–225.

70. Similä M, Kouki J, Martikainen P. 2003. Saproxylic beetles in managed and seminatural Scots pine forests: quality of dead wood matters. Forest Ecology and Management 174:365–381.

71. Sire L et al. 2022. Climate-induced forest dieback drives compositional changes in insect communities that are more pronounced for rare species. Communications Biology 5:1–17. Nature Publishing Group.

72. Stokland JN, Siitonen J, Jonsson BG. 2012. Biodiversity in Dead Wood. Cambridge University Press, Cambridge. Available from https://www.cambridge.org/core/books/biodiversity-in-dead-wood/32EA8DA79A503B95795384FFA5BC993D (accessed May 21, 2020).

73. Swanson ME, Franklin JF, Beschta RL, Crisafulli CM, DellaSala DA, Hutto RL, Lindenmayer DB, Swanson FJ. 2011. The forgotten stage of forest succession: early-successional ecosystems on forest sites. Frontiers in Ecology and the Environment 9:117–125.

74. Thorn S et al. 2018. Impacts of salvage logging on biodiversity: A meta-analysis. Journal of Applied Ecology 55:279–289.

75. Thorn S et al. 2020. Estimating retention benchmarks for salvage logging to protect biodiversity. Nature Communications 11:4762. Nature Publishing Group.

76. Thorn S, Bässler C, Bernhardt-Römermann M, Cadotte M, Heibl C, Schäfer H, Seibold S, Müller J. 2016. Changes in the dominant assembly mechanism drive species loss caused by declining resources. Ecology Letters 19:163–170.

77. Thorn S, Bässler C, Svoboda M, Müller J. 2017. Effects of natural disturbances and salvage logging on biodiversity – Lessons from the Bohemian Forest. Forest Ecology and Management 388:113–119.

78. Turner MG. 2010. Disturbance and landscape dynamics in a changing world. Ecology 91:2833–2849.

79. Turner MG, Seidl R. 2023. Novel Disturbance Regimes and Ecological Responses. Annual Review of Ecology, Evolution, and Systematics 54:null.

80. Viljur M-L et al. 2022. The effect of natural disturbances on forest biodiversity: an ecological synthesis. Biological Reviews 97:1930–1947.

81. Wermelinger B. 2021. Forest Insects in Europe: Diversity, Functions and Importance. CRC Press, Boca Raton.

82. Wermelinger B, Duelli P, Obrist MK. 2002. Dynamics of saproxylic beetles (Coleoptera) in windthrow areas in alpine spruce forests. Forest Snow and Landscape Research 77.

83. Wermelinger B, Obrist MK, Duelli P, Schneider Mathis D, Gossner MM. 2025. Two decades of arthropod biodiversity after windthrow show different dynamics of functional groups. Journal of Applied Ecology 62:371–387.

84. Weslien J, Djupström LB, Schroeder M, Widenfalk O. 2011. Long-term priority effects among insects and fungi colonizing decaying wood. Journal of Animal Ecology 80:1155–1162.

85. Zemlerová V et al. 2023. Natural Disturbances are Essential Determinants of Tree-Related Microhabitat Availability in Temperate Forests. Ecosystems DOI: 10.1007/s10021-023-00830-8. Available from 10.1007/s10021-023-00830-8 (accessed March 30, 2023).

