## Supplementary material for "From forest decline to salvage logging: cascading impacts on saproxylic beetle diversity"

### Supplementary material of the article "From forest decline to salvage logging: cascading impacts on saproxylic beetle diversity"

Jérémy Cours [1,2,3,4], Aurélien Sallé [5], Laurent Larrieu [6,7], Carlos Lopez-Vaamonde [4,8], Jörg Müller [9,10], Guilhem Parmain [1], Simon Thorn [11,12,13], Alain Roques [4], Christophe Bouget [1]

**Keywords**: forest dieback, forest die-off, temperate forest, lowland forest, highland forest, montane forest, biodiversity, climate change, drought, pest insect outbreak

[1] UR EFNO, INRAE, Domaine des Barres, Nogent-sur-Vernisson, France

[2] Department of Biological and Environmental Science, University of Jyväskylä, Jyväskylä, Finland

[3] School of Resource Wisdom, University of Jyväskylä, Jyväskylä, Finland

[4] INRAE, URZF, Orléans, France

[5] Laboratory of Physiology, Ecology and Environment, University of Orléans, INRAE, Orléans, France

[6] INRAE, UMR DYNAFOR, University of Toulouse, Castanet-Tolosan, France

[7] CNPF-CRPF Occitanie, Tarbes, France

[8] IRBI, UMR 7261, CNRS-Université de Tours, Tours, France

[9] Bavarian Forest National Park, Freyunger Str. 2, 94481 Grafenau, Germany

[10] University of Würzburg, Department of Animal Ecology and Tropical Biology, Chair of Conservation Biology and Forest Ecology, Glashüttenstraße 5, 96181 Rauhenebrach, Germany

[11] Department of Biology, Philipps Universität Marburg, Marburg, Germany

[12] Hessian Agency for Nature Conservation, Environment and Geology, Biodiversity Center, Giessen, Germany

[13] Czech Academy of Sciences, Biology Centre, Institute of Entomology, České Budějovice, Czech Republic

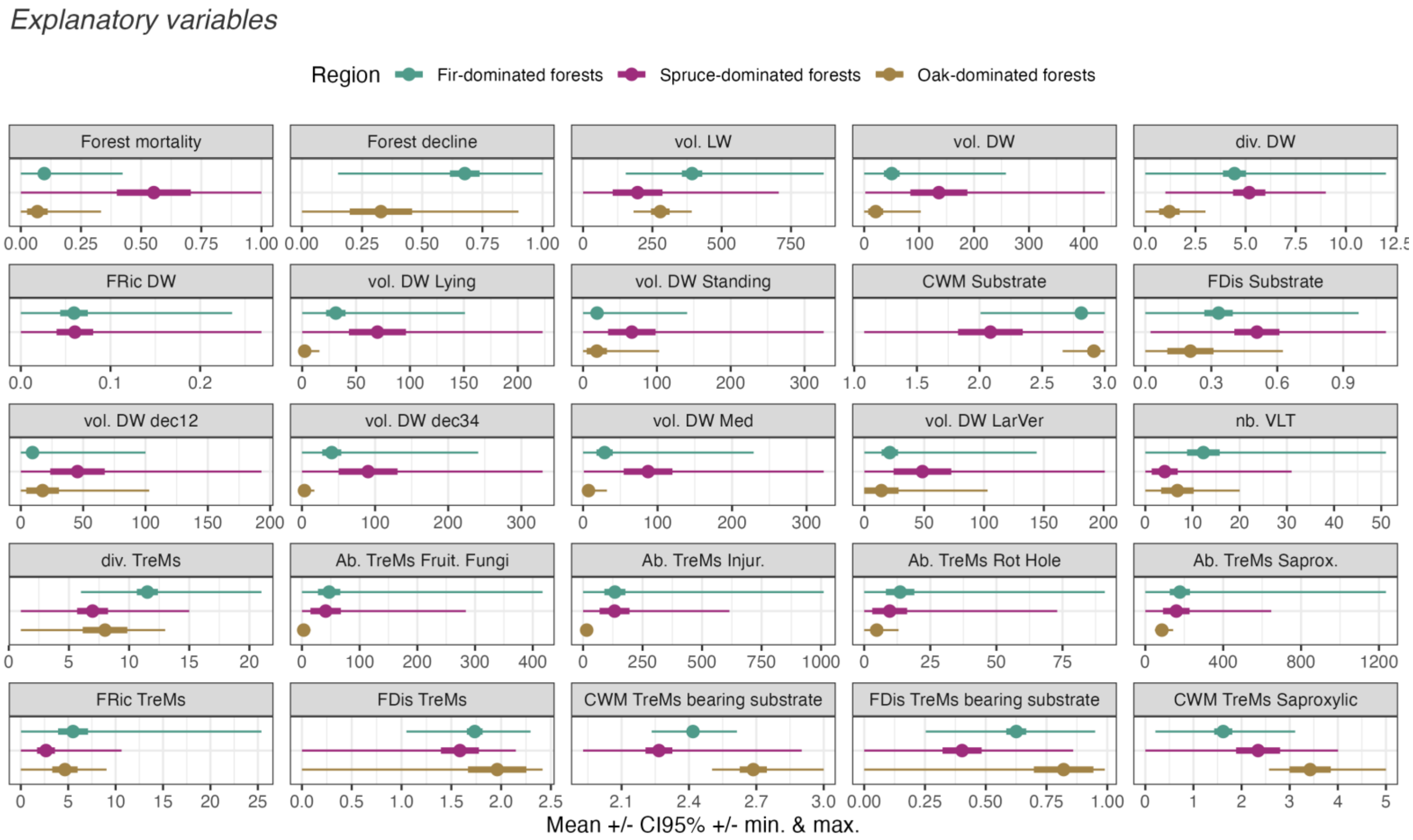


**Figure S1.** Mean and range values (bold line = confidence interval at 95%, thin line = range of the minimum and maximum values) of the explanatory variables. Deadwood volume is presented in $m^3.ha^{-1}$. The abundance of TreMs is to understand as the number of trees bearing the specific set of TreMs, in $nb.ha^{-1}$.

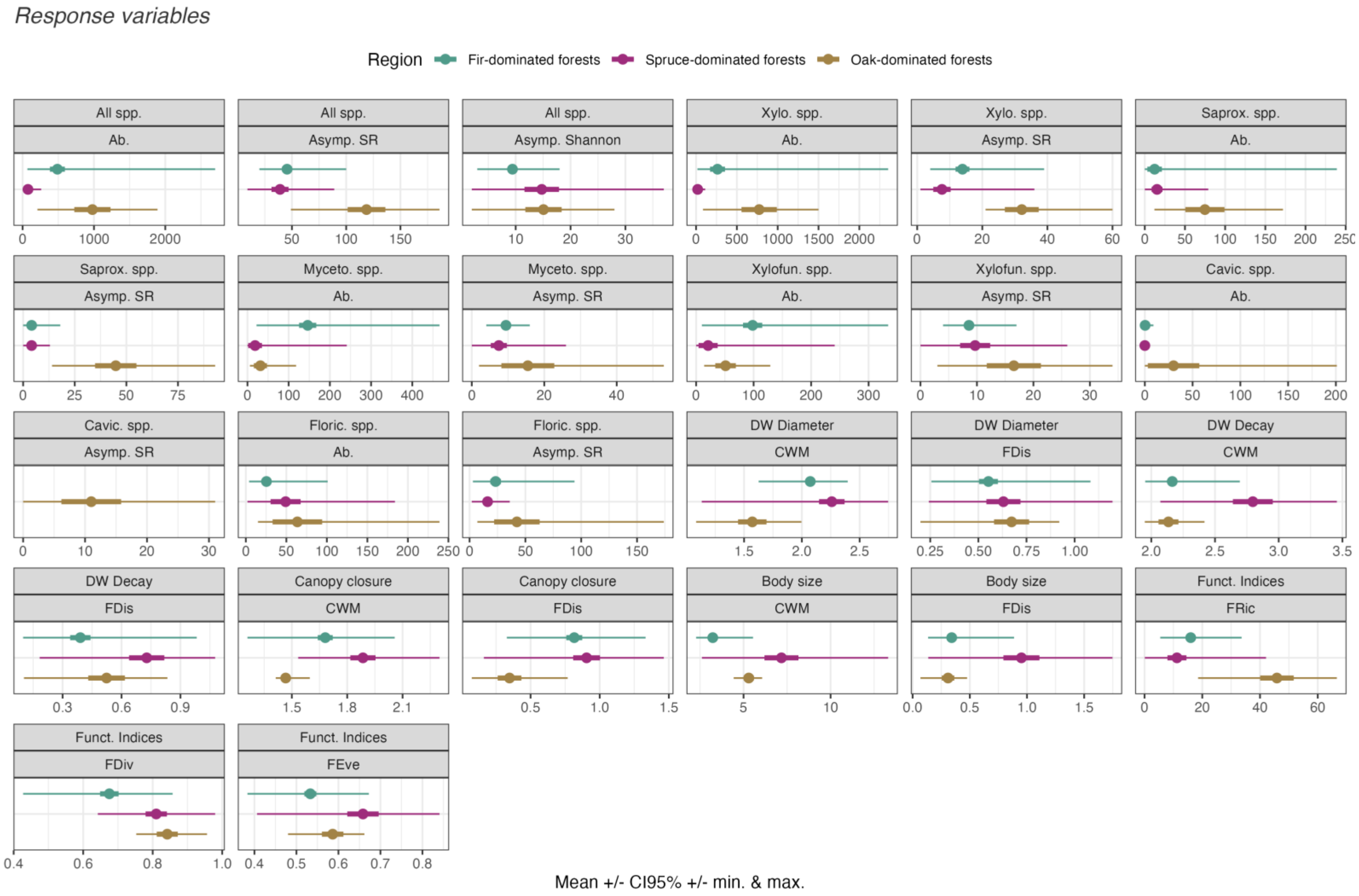


**Figure S2.** Mean and range values (bold line = confidence interval at 95%, thin line = range of the minimum and maximum values) of the response variables.

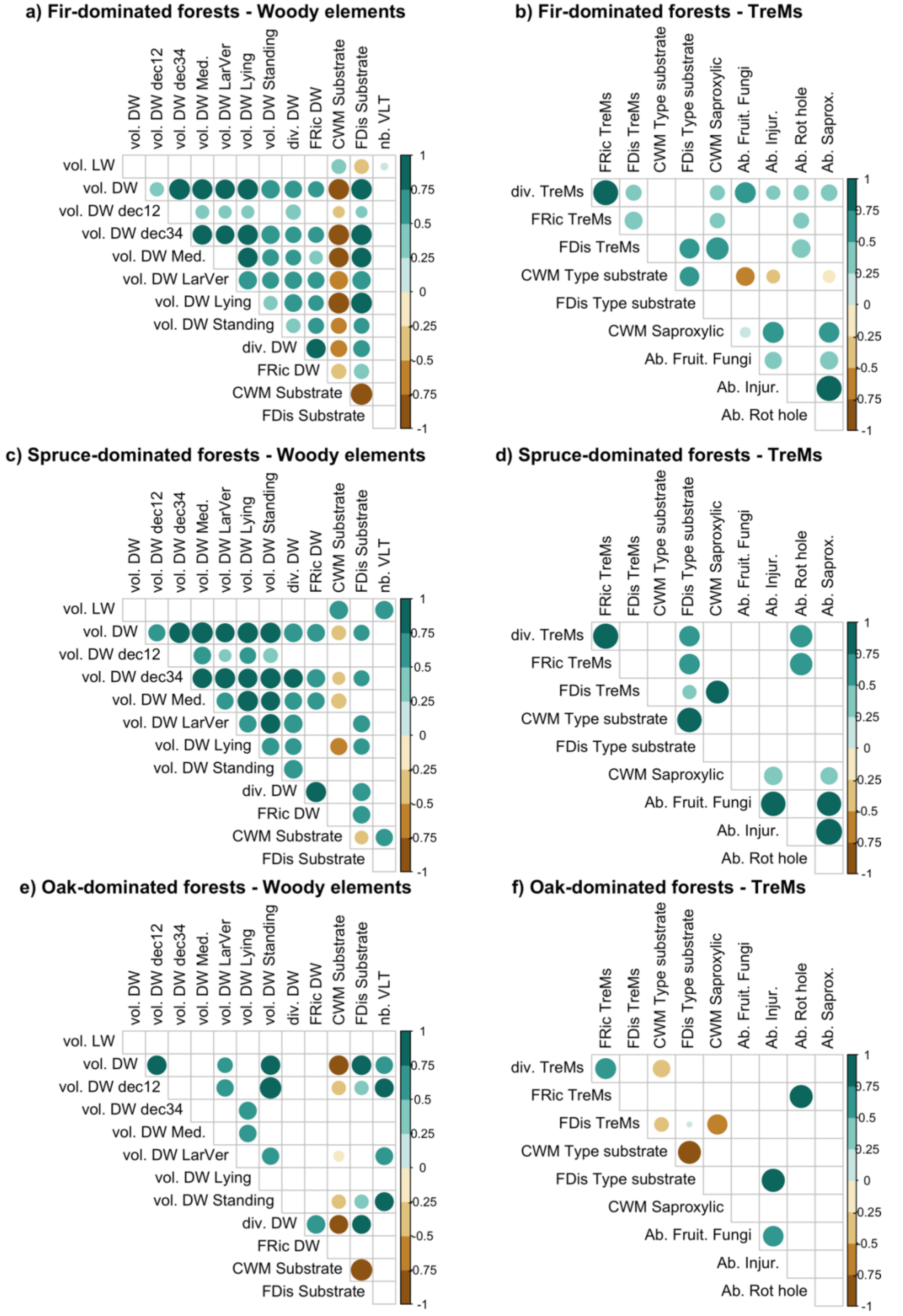


**Figure S3.** Spearman's correlation between the predictors (left: woody elements, right: TreMs) tested in SEMs, for each context. Only the significant correlations are shown (at $p < 0.05$).

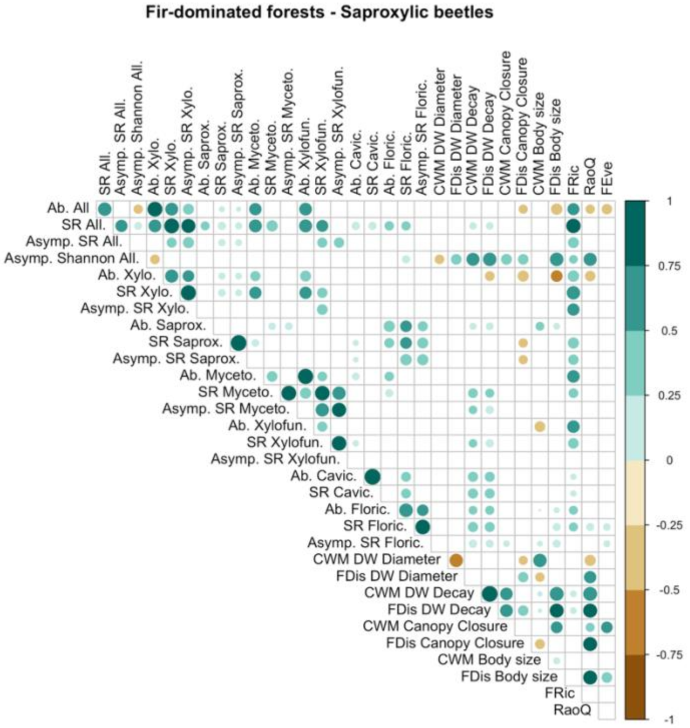


**Figure S4.** Spearman's correlation between biodiversity variables in fir-dominated forests.

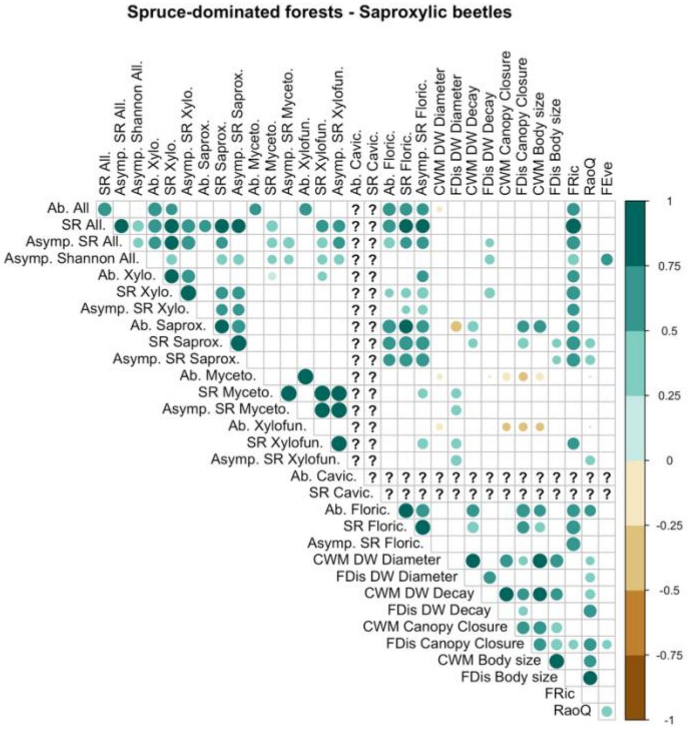


**Figure S5.** Spearman's correlation between biodiversity variables in spruce-dominated forests.

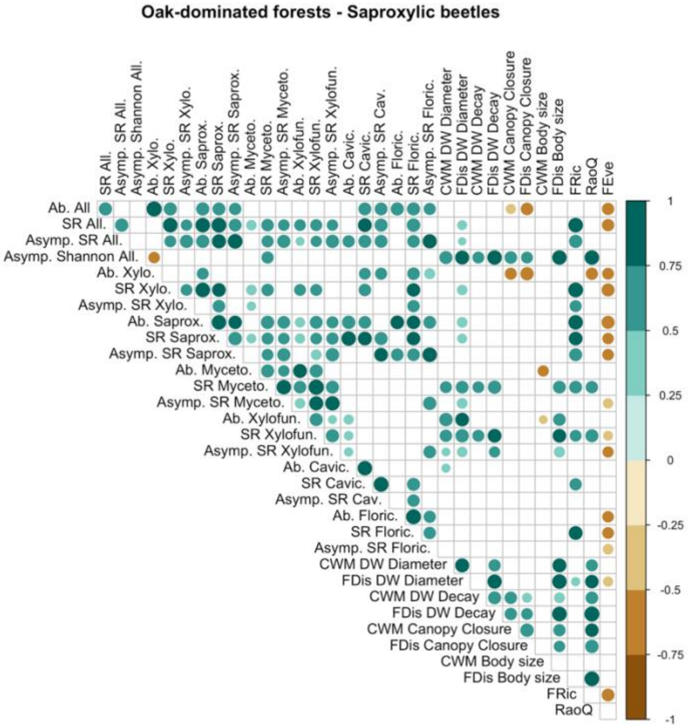


**Figure S6.** Spearman's correlation between biodiversity variables in oak-dominated forests.

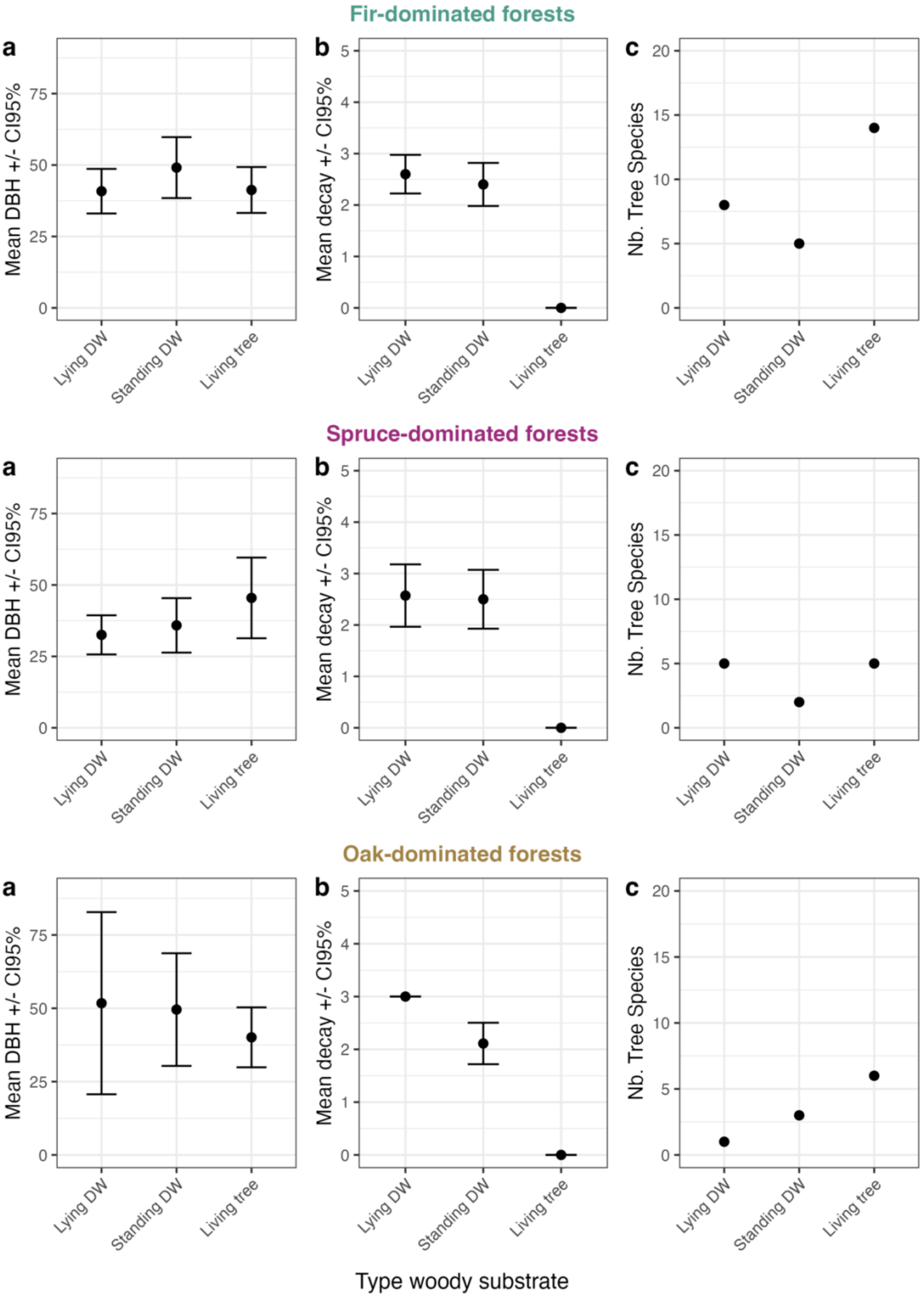


**Figure S7.** Characteristics of woody substrates in each forest context. Average values and confidence interval based on substrate trait data.

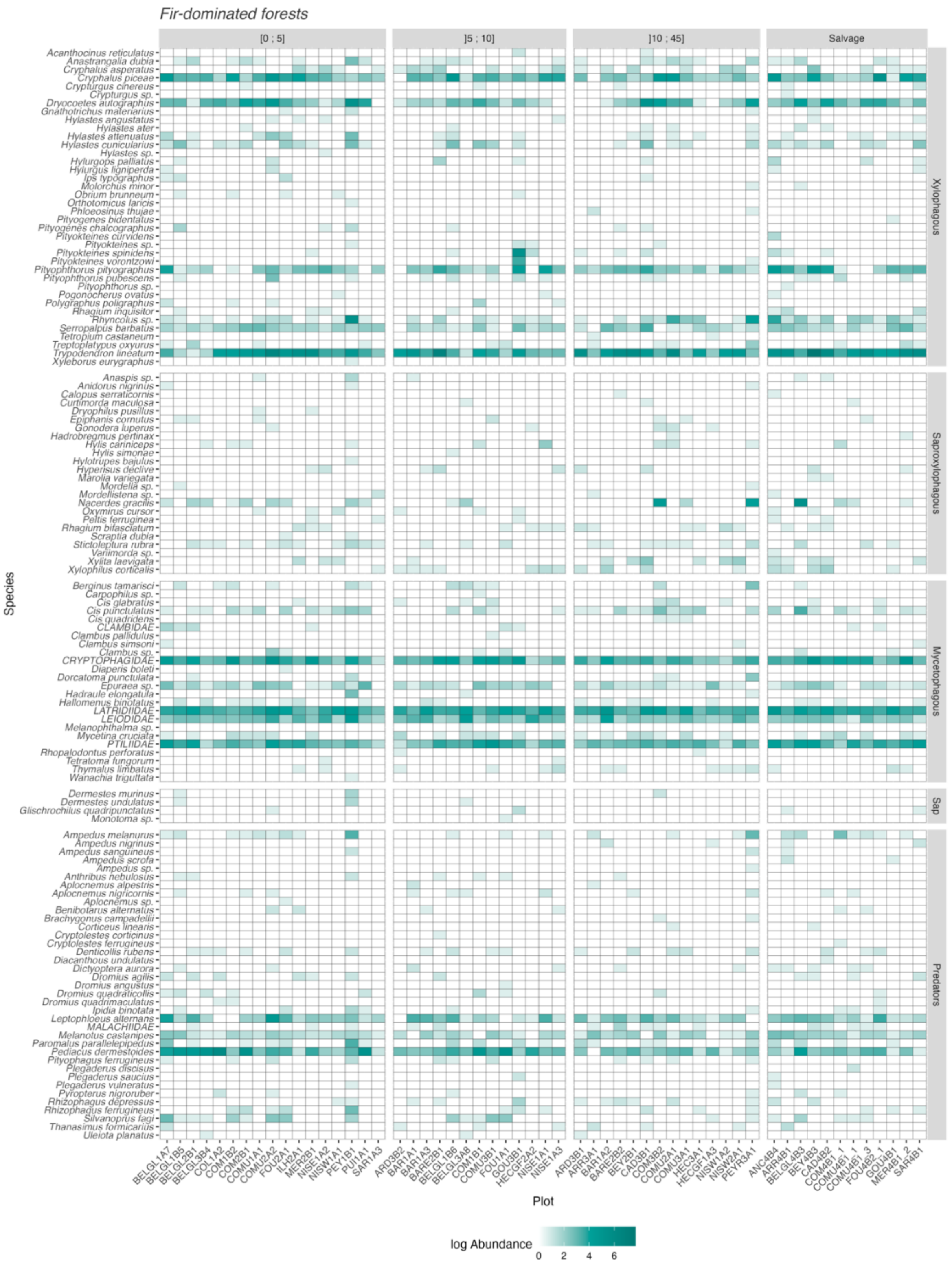


**Figure S8.** List of species in fir-dominated forests and their abundance (log-transformed) within each studied plot. Classes of forest mortality.

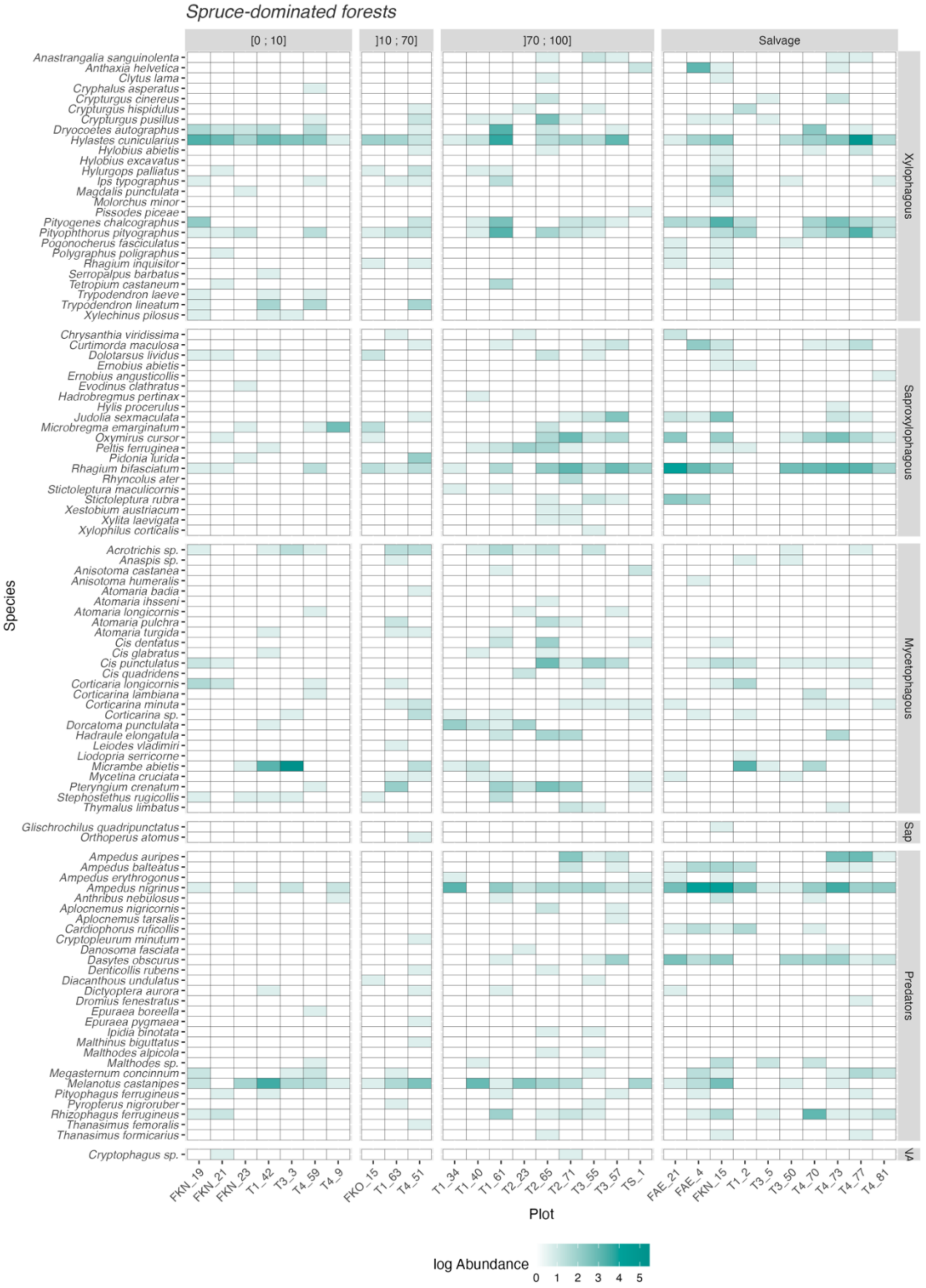


**Figure S9.** List of species in spruce-dominated forests and their abundance (log-transformed) within each studied plot. Classes of forest mortality.

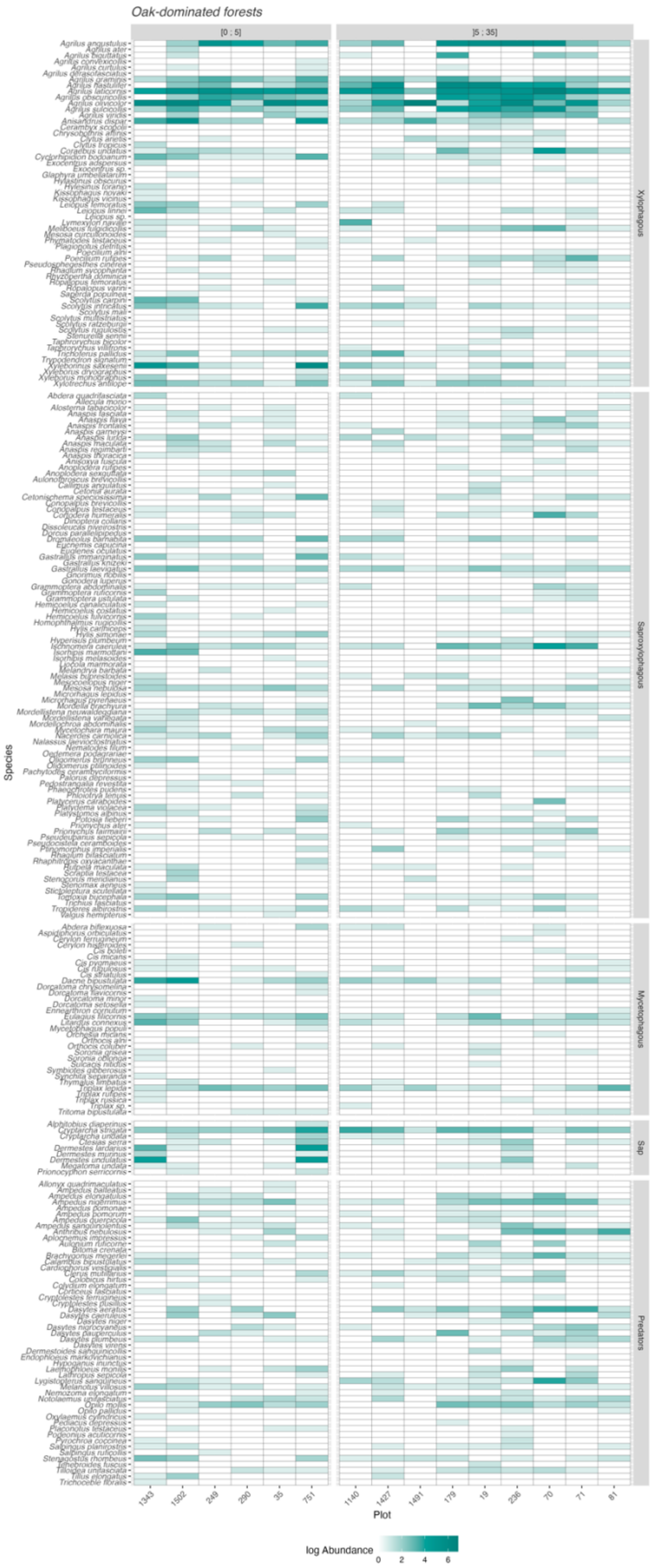


**Figure S10.** List of species in oak-dominated forests and their abundance (log-transformed) within each studied plot. Classes of forest mortality.
